# Kinematics of wings from *Caudipteryx* to modern birds

**DOI:** 10.1101/393686

**Authors:** Yaser Saffar Talori, Jing-Shan Zhao, Jingmai Kathleen O’Connor

## Abstract

This study seeks to better quantify the parameters that drove the evolution of flight from nonvolant winged dinosaurs to modern birds. In order to explore this issue, we used fossil data to model the feathered forelimb of *Caudipteryx*, the most basal non-volant maniraptoran dinosaur with elongate pennaceous feathers that could be described as forming proto-wings. In order to quantify the limiting flight factors, we created three hypothetical wing profiles for *Caudipteryx* representing incrementally larger wingspans, which we compared to the actual wing morphology as what revealed through fossils. These four models were analyzed under varying air speed, wing beat amplitude, and wing beat frequency to determine lift, thrust potential and metabolic requirements. We tested these models using theoretical equations in order to mathematically describe the evolutionary changes observed during the evolution of modern birds from a winged terrestrial theropod like *Caudipteryx. Caudipteryx* could not fly, but this research indicates that with a large enough wing span *Caudipteryx*-like animal could have flown, the morphology of the shoulder girdle would not actually accommodate the necessary flapping angle and metabolic demands would be much too high to be functional. The results of these analyses mathematically confirm that during the evolution of energetically efficient powered flight in derived maniraptorans, body weight had to decrease and wing area/wing profile needed to increase together with the flapping angle and surface area for the attachment of the flight muscles. This study quantifies the morphological changes that we observe in the pennaraptoran fossil record in the overall decrease in body size in paravians, the increased wing surface area in *Archaeopteryx* relative to *Caudipteryx*, and changes observed in the morphology of the thoracic girdle, namely the orientation of the glenoid and the enlargement of the sternum.

## Introduction

The origin of birds has still been a theme of considerable scientific debate [1–7]. Currently it is nearly universally accepted that Aves belongs to the derived clade of theropod dinosaurs, the Maniraptora [8,9]. The oviraptorosaur *Caudipteryx* is a member of this clade and the basal-most maniraptoran with pennaceous feathers [10–12]. The longest of these feathers are located distally on the forelimb, strongly resembling the ‘wings’ of birds. However, the relative length of these feathers compared to those in volant birds and the brevity of the forelimb itself compared to the si/e of the body and length of the hindlimbs all point to a clearly terrestrial animal [13–15]. It is understood that winged forelimbs must have evolved first for some other purpose and were later exapted for flight. Hypotheses regarding their original function range from ornamentation, temperature regulation, or locomotion [16–21]. The feathers on the forelimbs and tail in *Caudipteryx* are straight with symmetrical vanes similar to the wing feathers in flightless birds. It’s certainly clear that *Caudipteryx* couldn’t fly [22–24]. However, as a fairly fast cursorial animal, the presence of feathered distal forelimbs must have had some effect on the locomotion of *Caudipteryx*. The aerodynamic effect of the basal-most known forelimbs with pennaceous feathers has the potential to shed light on the evolution of flying wings in Paraves (see Supplementary Materials for more basic information about the flight kinematics of bird’s wings).

We mathematically modeled *Caudipteryx* with three hypothetical wing sizes (morphotypes **B**, **C**, and **D**) across a range of input values for flapping angle, wing beat frequency and velocity (the hypothetical wingspans are based on such modern birds listed in S1 Table and the wing profiles are inspired by an eagle [25]). We explore what values would be necessary to allow *Caudipteryx* to fly. This analysis quantifies the physical constraints that exclude the cursorial *Caudipteryx* from engaging in volant behavior despite the presence of feathered forelimbs and hint at the evolutionary changes necessary to evolve flight in the maniraptoran lineage from a cursorial animal with small wings like *Caudipteryx* to the neornithine condition, which is supported through observations from the fossil record of pennaraptorans.

**Fig 1.**
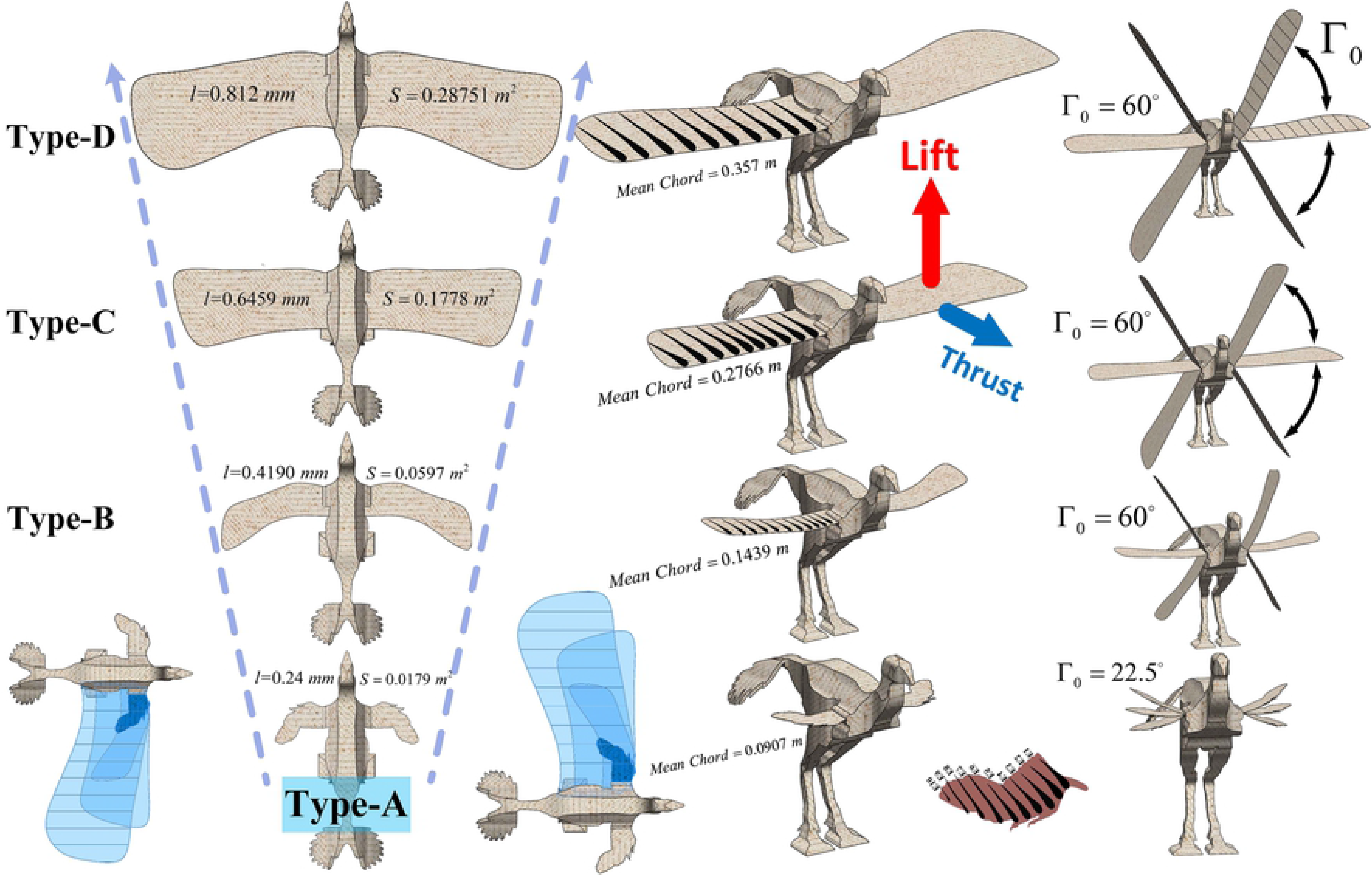
Type A is an actual *Caudipteryx’s* model and the wings in Types B, C and D are assumed in different sizes changing in linear form.

## Materials and methods

In order to achieve a reasonable accuracy in analyses, each wing type (Fig 1) is divided into ten elements, each modeled using unsteady aerodynamics in order to capture lift, thrust/drag and required power (Fig 2). Type-**A** is based on fossil data and represents a realistic estimate of wing length (***l***) which we measure to be 0.24 meters. Types-**B**, **C** and **D** are hypothetical models with increasing wing lengths of 0.419, 0.6459 and 0.812 meters, respectively, and for the sake of simplicity, we approximately calculate the areas of the wings with the equivalent rectangular ones. Wings size in terms of area ***S*** (***m***^2^), mean chord length ***C*** (***m***), aspect ratio (***λ***), flapping angle (Γ_0_) and other wing angles describe the state of the wing (Fig 1 and Table 1). The element’s motion includes a plunging velocity ***ḣ*** and pitch angle (twist angle) ***θ***. We assumed that the wing’s aspect ratio is large enough to pass flow through each element in the mean stream direction. Hence, the normal force ***dN*** of the element’s total attached flow, is equal to the normal force ***dN**_a_* of element’s circular plus apparent mass effect, as an additional normal force contribution which acts at the mid chord. Delaurier and Larijani have given the formulas below based on mathematics, tests and experiments [26–32].

**Table 1.** Specifications of Types-**A**, **B**, **C** and **D**

|  | Type-A | Type-B | Type-C | Type-D |
| --- | --- | --- | --- | --- |
| $l(\mathbf{m})$ | 0.24 | 0.419 | 0.6459 | 0.812 |
| $S(\mathbf{m}^2)$ | 0.0179 | 0.0597 | 0.1778 | 0.28751 |
| $C(\mathbf{m})$ | 0.0907 | 0.1439 | 0.2766 | 0.3575 |
| $\lambda$ | 3.2 | 2.94 | 2.35 | 2.3 |
| $\bar{\theta}_a(\text{deg})$ | 0 | 0 | 0 | 0 |
| $\bar{\theta}_w(\text{deg})$ | 15 | 20 | 20 | 20 |
| $\Gamma_0(\text{deg})$ | 22.5 | 60 | 60 | 60 |
| $\alpha_0(\text{deg})$ | 30 | 10 | 10 | 10 |
| $\beta_0(\text{deg})$ | 180 | 10 | 10 | 10 |
| $\eta_s$ | 0.9 | 0.9 | 0.9 | 0.9 |

**Fig 2.**
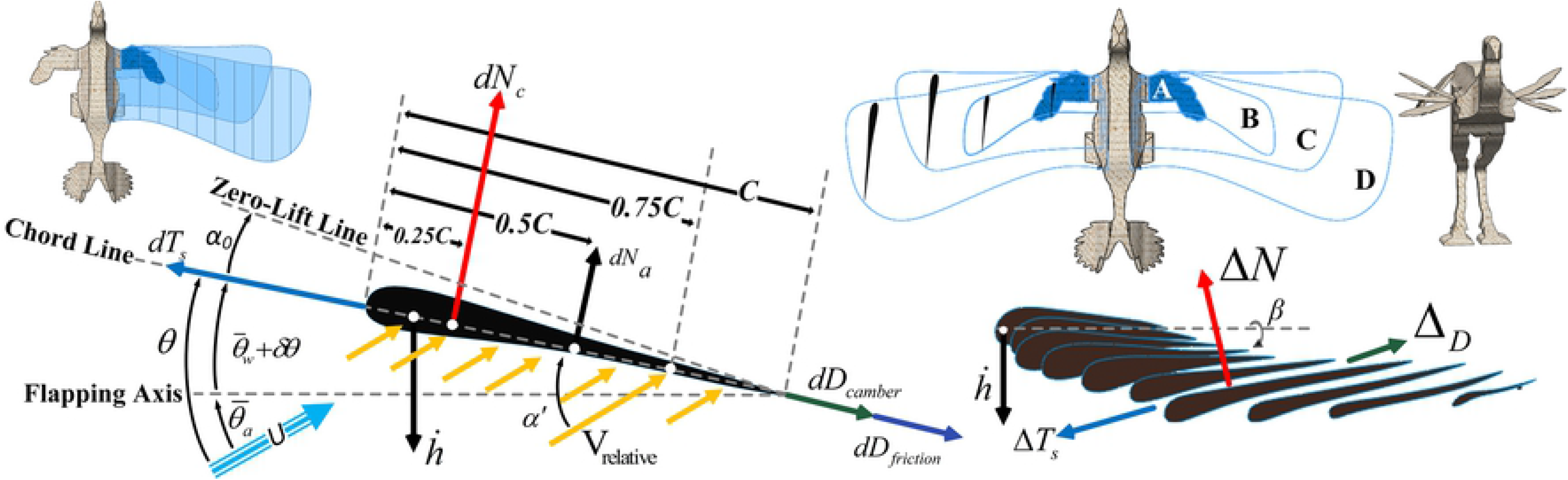
Kinematics model of flapping flight for each element of the wing of *Caudipteryx*.

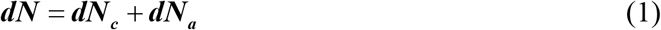

The normal force of the element’s circular and the apparent mass effect are defined as below

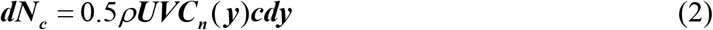

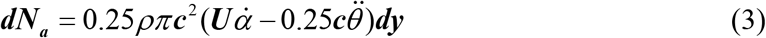

where *ρ* is the density of the airflow, *V* is the flow’s relative velocity at the ¼ chord, 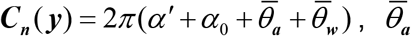 is the angle between flapping axis and mean stream velocity 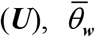 is the mean angle between chord and flapping axis, *α*_0_ is the angle of the zero lift line the value is fixed for airfoil in each situation along the wing), *α*’ is the flow’s relative angle of attack at the ¾ chord (Fig 2), it’s given by

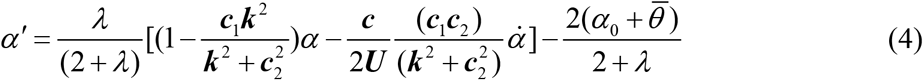

where *λ* is the aspect ratio of the wing of *Caudipteryx, c* is the wing element chord length, ***k*** = ***cω*** / **2*U*** which is the reduced frequency (*ω* is in radians and *ω* = 2*π**f***, and ***f*** is in hertz). Equation (4) is simplified by formulation of the modified Theodorsen function which was originally presented by Jones [33], *c*_1_ = 0.5*λ*/(2.32 + *λ*) and *c*_2_ = 0.181 + (0.772/*λ*). *α* and *α̇* are given by

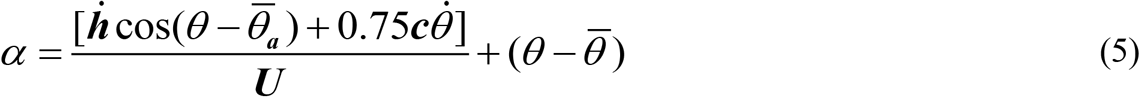

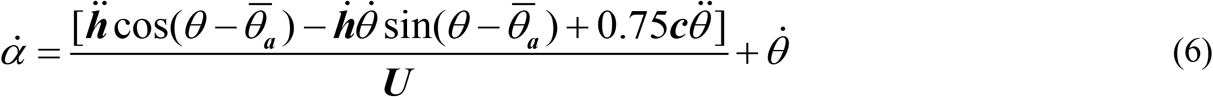

where *α* is the relative angle of attack at the ¾ chord. The pitch angle *θ* is 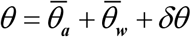 where *δθ* is the dynamically varying pitch angle (Fig 2). The *θ*(***y, t***) is a function of ***y*** and *ω*. Therefore, *δθ*(***y, t***) = −*β*_0_*y* sin(*ω**t***) which prescribes the twist of wing of *Caudipteryx* where *β*_0_ is a constant representing the twist angle per unit distance along the wing span (°/m). Hence, the first and second derivatives of *θ*(***y, t***) with respect to time are written as

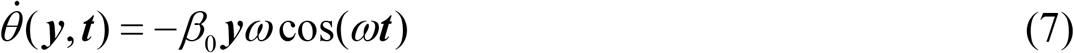

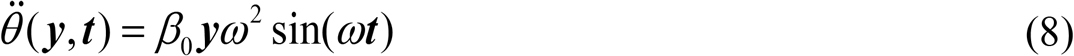

The plunging displacement of each element is ***h*(*t*)** = **(Γ_0_*y*)** × **cos(*ωt*)** (the imposed motion), where Γ_0_ is the maximum flapping amplitude and ***y*** is the distance between flapping axis (wing base) and center of a wing segment. Therefore, the plunging velocity ***ḣ*** (**t**) and acceleration ***ḧ*(*t*)** at the leading edge are given as

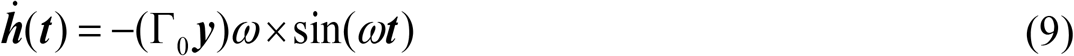

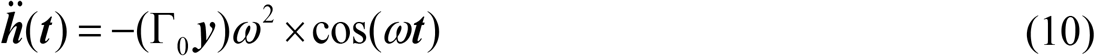

Pitch angle (twist angle) changes during flapping and these variations are proportional to flapping angle in a cycle (S2 Fig). The wing motion of *Caudipteryx* relative to ***U*** is included in the flow velocity ***V*** given in equation (2) and also term of *α’* is taken into account, ***V*** is

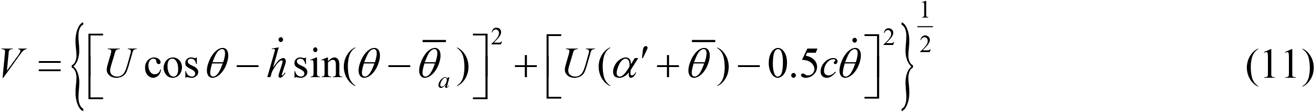

The element’s circulation distribution generates forces along the chord axis direction (Fig 2), hence the chordwise force due to the camber (***dD***_camber_) and chordwise friction drag due to viscosity are given by

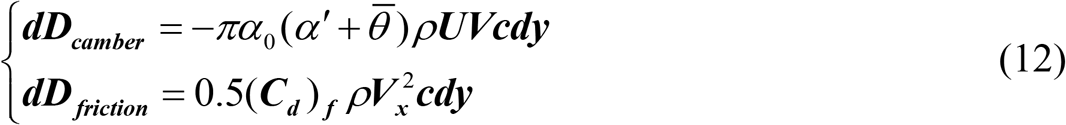

where (***C**_d_*)_f_ = 0.89/ (log(Re_*chord*_))^2.58^ is the friction drag coefficient [34] of the skin that is included in Reynolds’ number of local chord length and 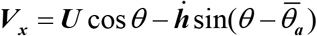 is the tangential flow speed to the element. For the two dimensional airfoil, Garrick expressed ***dT**_s_* for the leading edge as below [35,36]

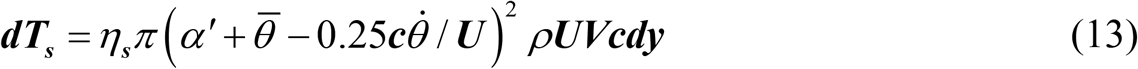

where *η_s_* is the efficiency term which presents that in reality because of viscous effects, the efficiency of leading edge of most airfoils are less than 100%. Therefore, the total chordwise force along the chord axis is expressed with

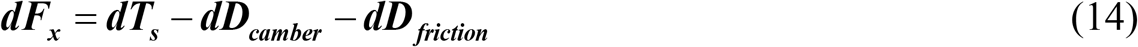

Therefore, the equations of element’s instantaneous lift and thrust are rewritten as

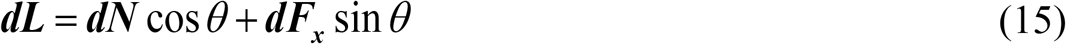

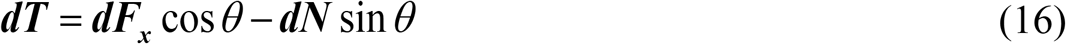

where ***dL*** and ***dT*** are the instantaneous lift and thrust of the element, respectively. To integrate along the wingspan and to get whole wing’s instantaneous lift and thrust, for wings, it is given by

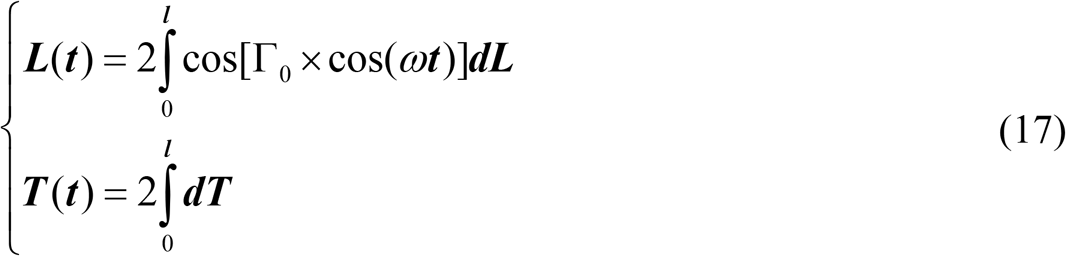

where *l* is the wing span length. The average of the lift and thrust can be obtained via integrating equation (17) over a cycle as *φ* = *ω**t***.

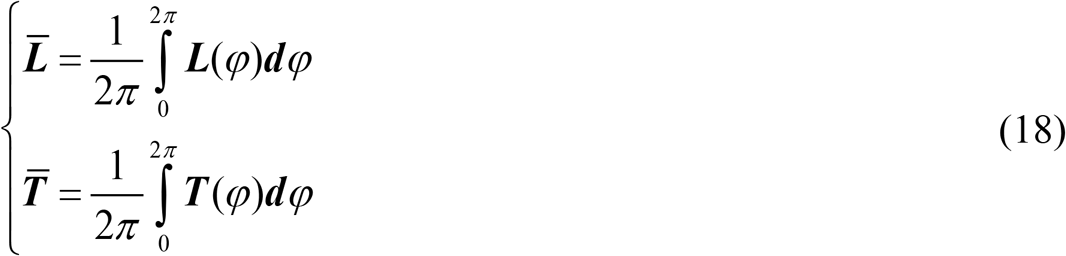

where 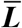 and 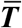 are the averages of the lift and thrust, individually. To calculate the required power against the forces, it is represented for attached flow by

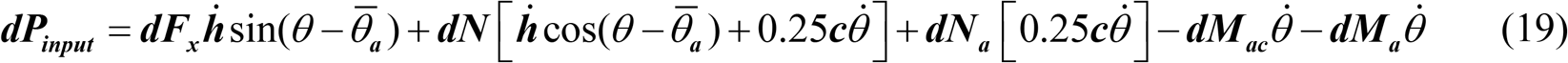

where ***dM**_ac_* is the element’s pitching moment about its aerodynamic center and ***dM**_a_* is composed of apparent chamber and apparent inertia moments are given respectively as follows

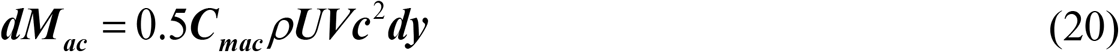

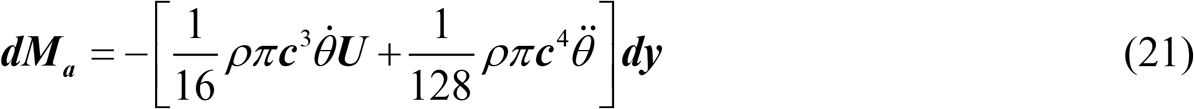

where **C*_mac_* is the coefficient of airfoil moment about its aerodynamic center. The instantaneously required power, ***P**_input_* (***t***), for a whole wing is derived below

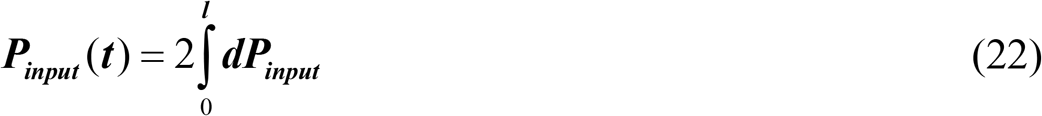

Then the average input power in a cycle is found from

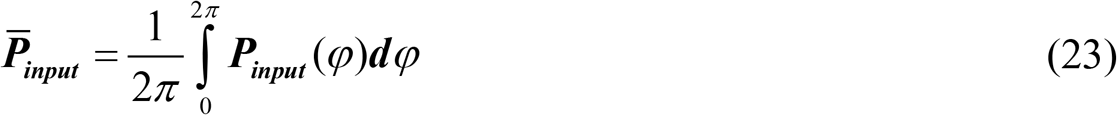

**Fig 3.**
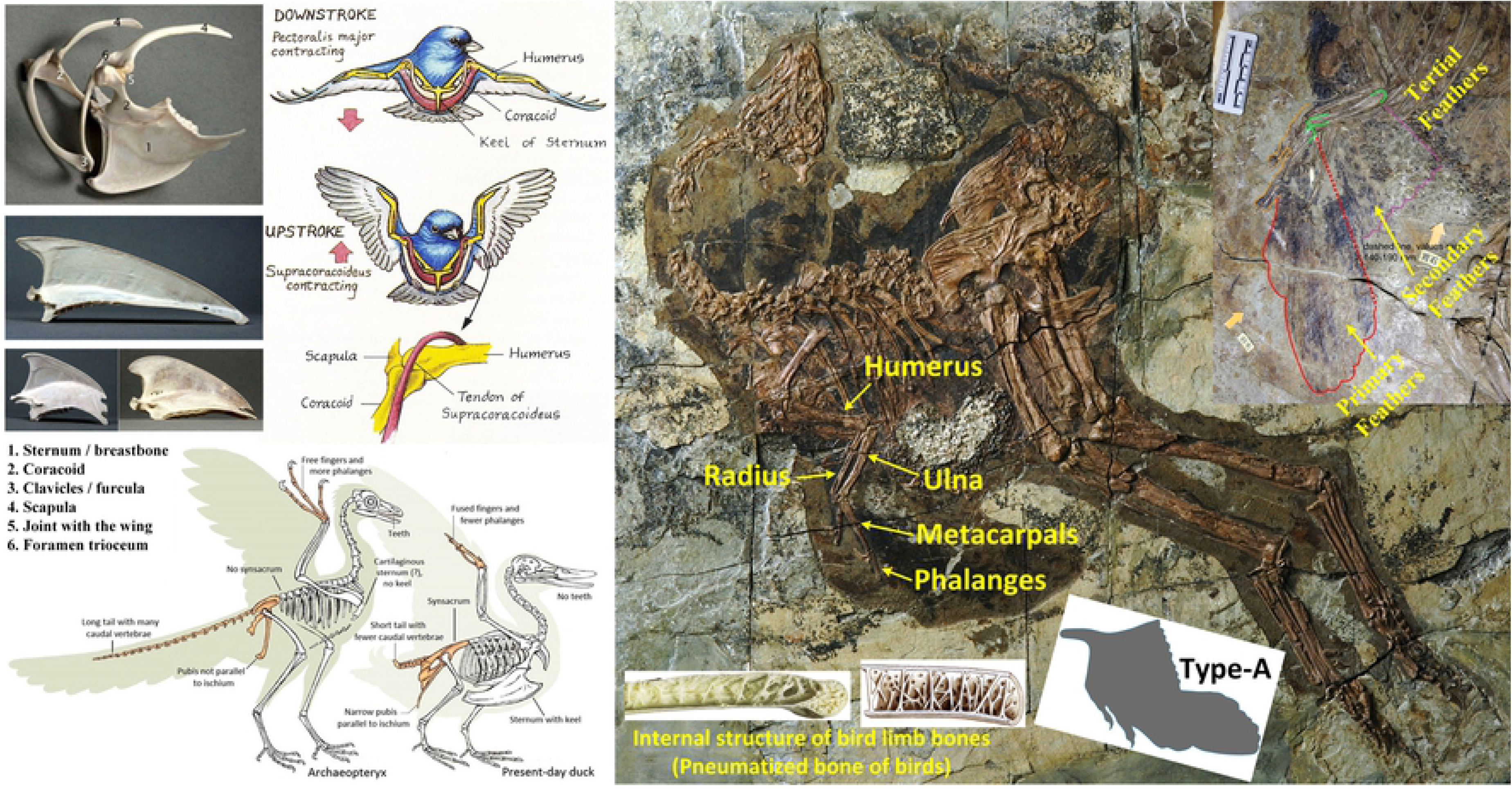
Illustrations of the avian flight apparatus in *Archaeopteryx* and living birds. A keeled sternum indicates increased capacity for flight based on the increased surface area for the attachment of the flight muscles. The realistic mode and reconstructed wing type (type-**A**) is based on *Caudipteryx* fossil [24].

The weight of body is computed about 50 N for the holotype of *Caudipteryx* dongi-IVPP V12344-Based on equation relating femur length to mass [37], the average of the lengths of the left and right femur are both 149 mm. With the formulas above, we compare the lift, thrust and power of each model with regards to the changes in wing angles (flapping angle (S4 Table), 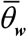 (S5 Fig) and dynamic twist angle (S6 Fig)), flapping frequency from lower frequency (1 Hertz) (S9 Fig and S10 Fig) to higher frequency (8 Hertz) (Fig 4) and velocity while the range of the air velocity changes from almost zero (U=0.05 m/s) (Fig 4, S8 Fig and S9 Fig) to higher speed (U= 10 m/s) (Fig 4 and S9 Fig). In the analyses on type-**A**, the wing angles are 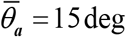, 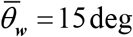, *α*_0_ = 30deg, *β*_0_ = 180deg deg and Γ_0_ = 22.5deg and in hypothetical models(types **B**, **C** and **D**), the corresponding parameters are 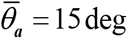, 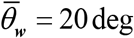, *α*_0_ = 1deg, *β*_0_ = 10 deg and Γ_0_ = 60 deg.

**Fig 4.**
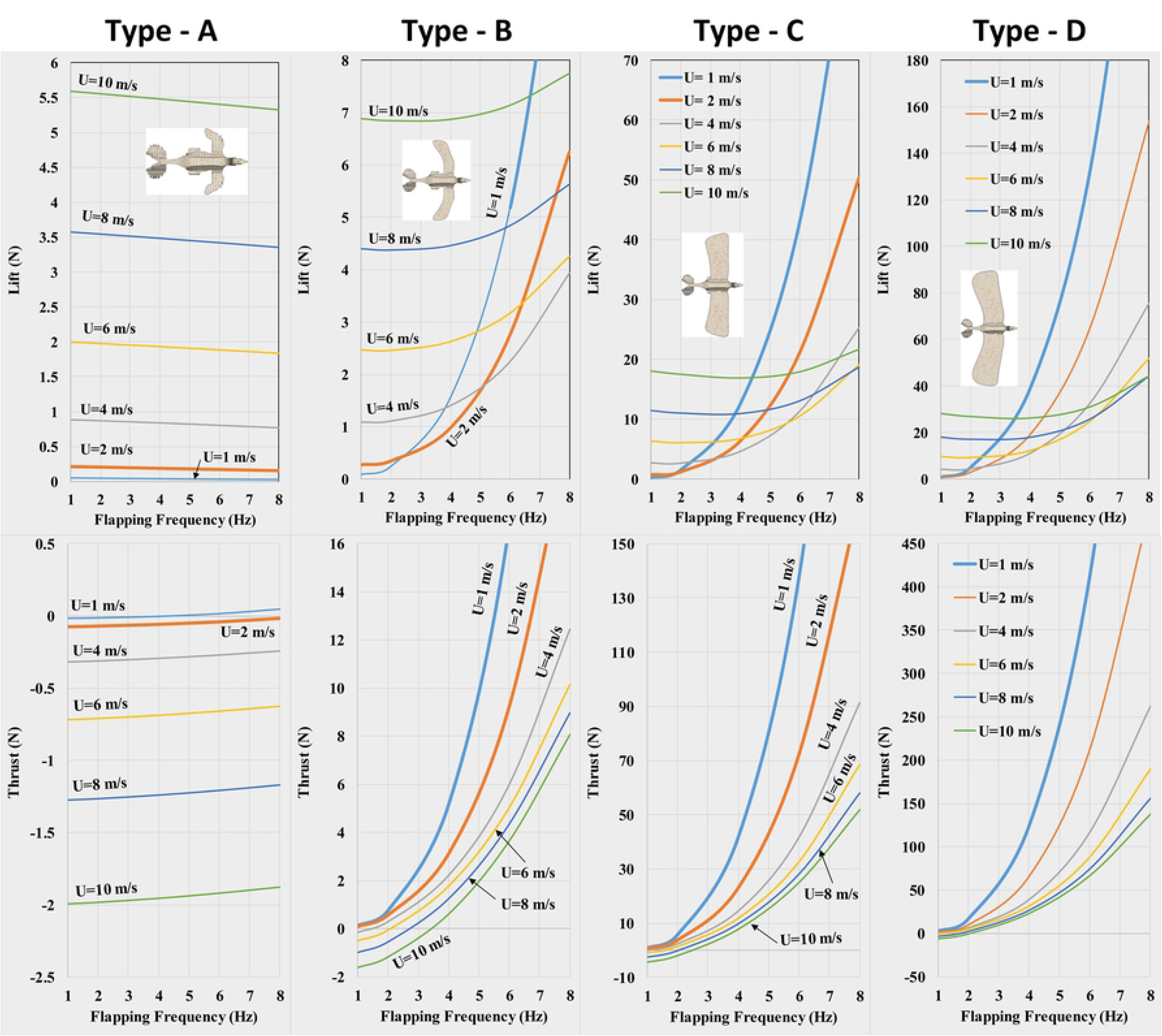
Variation in lift and thrust of *Caudipteryx* with respect to flapping frequency at any velocity.

## Results

When 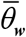 is 15 deg the realistic *Caudipteryx* (type-**A**) is capable of producing maximum thrust force but to have only vertical motion (maximum lift force with a small value of thrust), 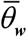 must be increased to 45 deg while *U* is 0.05 m/s (velocity is almost zero) (S5 Fig). In other hypothetical models (types **B**, **C** and **D**) the situation is almost similar to the realistic one (i.e. with the angle of 20 deg for maximum thrust and 45 deg for maximum lift). Hence, lift increases and thrust decreases as the angle of 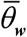 increases. In all models consumed power is proportional to the thrust force (S5 Fig) and the maximum thrust force requires maximum power.

The feathers in type-**A** were modeled as softer than those in modern birds (being primitive and symmetrical), and as the dynamic twist angle *β*_0_ expresses the stiffness of feathers along the wing span, we presumed *β*_0_ = 180° when lift and thrust are at their maximum values. The best and optimum value of this angle for three hypothetical models (types **B**, **C** and **D**) is 10 deg when the velocity is almost zero in one cycle (*f*=1Hz) (S6 Fig). In all models, as the flapping angle (wing beat amplitude) Γ_0_ increases, lift, thrust and the required energy also increase simultaneously (S4 Fig).

When birds supinate their wings, drag forces help the birds to brake. Larger wings moving at higher velocities produce greater braking forces (S7 Fig). Based on our analysis, the wings of *Caudipteryx* (type-**A**) utilizing both lower and higher wing beat frequencies at any velocity, are always capable of generating positive lift, but thrust force is only positive when the velocity is near zero and is negative during movement at any larger velocity (Fig 4). Hence in type-**A**, when flapping frequency and velocity increase, lift also increases but thrust converts to the drag force (negative thrust) causing a corresponding increase in metabolic demands (S3 Fig). In all hypothetical wings (types-**B, C** and **D**) lift is positive; an increase in flapping frequency at any velocity generates an increase in lift (Fig 4 and S9 Fig).

Although hypothetical wing type-B cannot provide sufficient lift force to overcome gravity (Fig 4 and S8 Fig), type-**C** is theoretically capable of supporting flight at lower velocity and higher wing beat frequency (namely U=1~2 m/s and frequency=6~8 Hz) but cannot generate large enough lift at higher velocities (Fig 4 and S9 Fig). The larger wings in type-**D** are capable of sustaining faster flight (8 m/s) with higher flapping frequency (*f*=4~8 Hz). Heavier birds flying at higher velocities require greater wing span. The obtained lift and thrust forces from Type-**A** deduced by experiments (S11 Fig) support our theoretical calculations (S2 Table).

## Discussion

Powered flight is the most physically demanding form of locomotion. Thus, during the course of evolution of flight from non-volant dinosaurs such as *Caudipteryx* to Aves, major skeletal adaptations needed to evolve in order to allow the necessary range of motion, accommodate the necessary musculature, and produce these adaptations in a light weight framework. These adaptations include modification of the glenoid to facilitate greater flapping angles and enlargement of the sternum to support the increased musculature necessary to generate significant thrust forces. This had to be paired with the ability to meet the metabolic demands of sustaining flight (the neornithine digestive system is highly specialized) [38–40]. We can infer these changes based on observations of the fossil record. However, the aerodynamic limitations that drove these evolutionary changes have never been explored.

In order to explore this idea, we tested the impact of increasing wing size in *Caudipteryx* within its actual skeletal framework. We considered three hypothetical wing morphologies (types **B, C**, and **D**) that vary in wing span, wing profile, wing chord, and aspect ratio (Fig 1). The goal in this study is to assess the effects of increasing wing size in a terrestrial form in order to identify which variables are limiting volant behavior. We focused on flight kinematic parameters during take off. The flight kinematics formulas utilized here to calculate lift, thrust and power allow us to compare all states in varying frequencies, velocities and flapping angles while ranging air stream velocity changes from almost zero to high speeds. This provides a comprehensive understanding of all important flight parameters.

In type **A**, the feathers are assumed to be weaker than those in types **B, C** and **D** and the flapping angle is much lower, which reflects the basal position of these non-aerodynamic feathers and the morphology of the glenoid. Increasing frequency and flapping angle from ±22.5 deg to ±60 deg (in types **B, C** and **D**) generates higher lift force (see S3 Table and S4 Table in Supplementary Materials for more information about the values of flapping beats and flapping angles). However, the theoretical requirements for metabolic power and muscle performance are unrealistic. This quantifies the observed changes in the scapulocoracoid observed during the evolution of birds from non-volant dinosaurs. Our analyses express that during low flapping frequency (**f**=3 Hz) and when the wind velocity is almost zero in wing types **B** and **C** (near type-**C**) the lift force generated by the wings is nearly enough to overcome gravity for take off. At low velocity and with a higher hindlimb frequency, type-**C** is able to meet the lift requirements. In this type, although the wings would not be able to provide enough force to fly faster, type **C** can supply the capability of flying at lower speed. The wings in type **D** are larger allowing the model *Caudipteryx* to fly faster than 8 m/s with higher wing beat frequency. Not surprisingly, the largest wings (type **D**) generate the greatest lift, thrust forces, and drag force during braking. However, as flapping frequency increases, so do the required thrust forces and metabolic requirements.

The aerodynamic behavior of the realistic *Caudipteryx* model, reconstructed from fossil material (type **A**), reveals that the wings could generate lift but at any velocity produced more drag than thrust, and thus probably did not evolve in this taxon for aerodynamic function. The analyses indicate that for a given lift or thrust force, all hypothetical types of *Caudipteryx* would have to increase wing beat frequency to increase their velocities. It depends on wingspan that produces capability of flying, those generations of the new birds whose wing’s profile is similar to those of types **B** and **C**, they need to decrease their self-weight during evolution to obtain enough lift and thrust for flight in accordance with the weight. Heavy modern birds (such as Black-browed albatross or Wandering albatross-see Table S1 in Supplementary Materials) shall have large enough wingspan more than that of type **C** to keep balance between lift force and their body weight.

In order to take off to overcome gravity, sustain flapping flight and maintain lift forces to oppose body-weight, flapping frequency, and flapping angle (as the most significant parameter) and wing span had to increase. At small flapping angles, like those present in *Caudipteryx* and other non-volant maniraptorans, the wing beat cannot generate large enough lift and thrust. This verifies the observed changes in shoulder architecture in the lineage to Aves from a more ventrolateral (non-volants) or laterally facing (*Archaeopteryx*) glenoid to the dorsolaterally facing glenoid in birds–clearly the wings of *Caudipteryx* are the smallest among the wings of known Mesozoic pennaraptorans and the birds, and the wings of volant dromaeosaurid *Microraptor* are much larger. Heavier flying birds need more lift to become airborne and thus require larger wingspans with large flapping angles and strong flight muscles.

Depending on the wing profile, the flapping angle should be more than 60 degrees (i.e. with a total range of motion of 120 degrees). To accommodate this range of motion, it required the dorsal expansion of the glenoid and an increase in wing musculature to manipulate the larger wings against greater aerodynamic resistance. This would drive up the metabolic cost of flight so high as to make this form of locomotion inefficient. However, this can be obviated by reducing the cost of flapping flight with an overall reduction in body weight.

Observations from fossils suggest that this was achieved both through a reduction in overall body size and by the evolution of additional pneumaticity and thinner bone cortices. Morphological changes of the skeleton such as the reduction of tail, and modification of numerous biological systems such as the loss of the right ovary and evolution of a highly efficient digestive system. This ultimately produced a body shape that was well adapted to spindle shaped, generating less resistance (drag force) during flight (Fig 3) [25,40,41]. Therefore, it is important for efficient flight to decrease body mass and to increase musculature, wing profile, aspect ratio, flapping angle in the evolution from non-volant maniraptoran dinosaur to modern birds.

## Acknowledgements

The authors acknowledge the kind suggestions from Prof. Dr. Corwin Sullivan from the Department of Biological Sciences, University of Alberta, Canada. Prof. Dr. Zhong-He Zhou and Prof. Dr. Min Wang from the Key Laboratory of Vertebrate Evolution and Human Origins, Institute of Vertebrate Paleontology and Paleoanthropology, Chinese Academy of Sciences, Beijing, 100044, P. R. China. This project was supported by the National Natural Science Foundation of China under grant 51575291, the National Major Science and Technology Project of China under grant 2015ZX04002101, State Key Laboratory of Tribology, Tsinghua University, and the 221 program of Tsinghua University.

## Author Contributions

Author contributions: Y. S. deduced formulas and prepared programs, simulations, tables and Figures and wrote the first draft of the manuscript; Y. S. and Y. F. accomplished the experiments and completed the 3D reconstruction and video of the *Caudipteryx* robot; J.-S. supervised the project and proposed the experiment principle; Y.-S., Y. F., J.-S. and J. K. completed the Cladogram of *Caudipteryx* and investigation of the feathers of the dinosaurs; J. K. provided the fossil and analysis of dinosaurs and provided the major suggestions in revision; All authors discussed the results and commented on the manuscript and contributed ideas to manuscript development and data analysis.

## Supporting information

### S1 Text. Supplementary materials

**S1 Fig.**
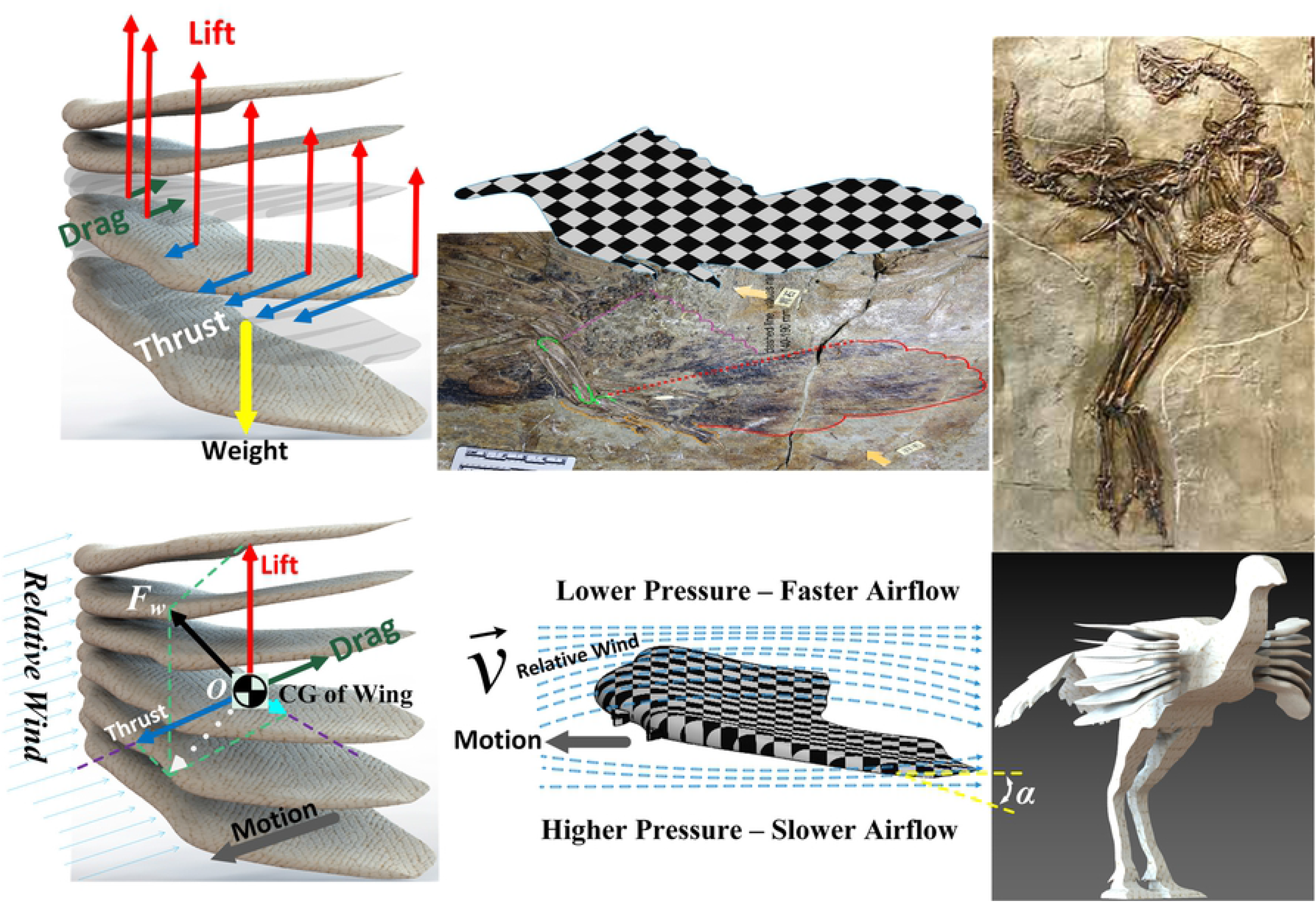
Reconstructed *Caudipteryx* via software in accordance with the fossil. Bernoulli effect and lift, thrust, weight and drag loads are represented along the wing span [2].

**S2 Fig.**
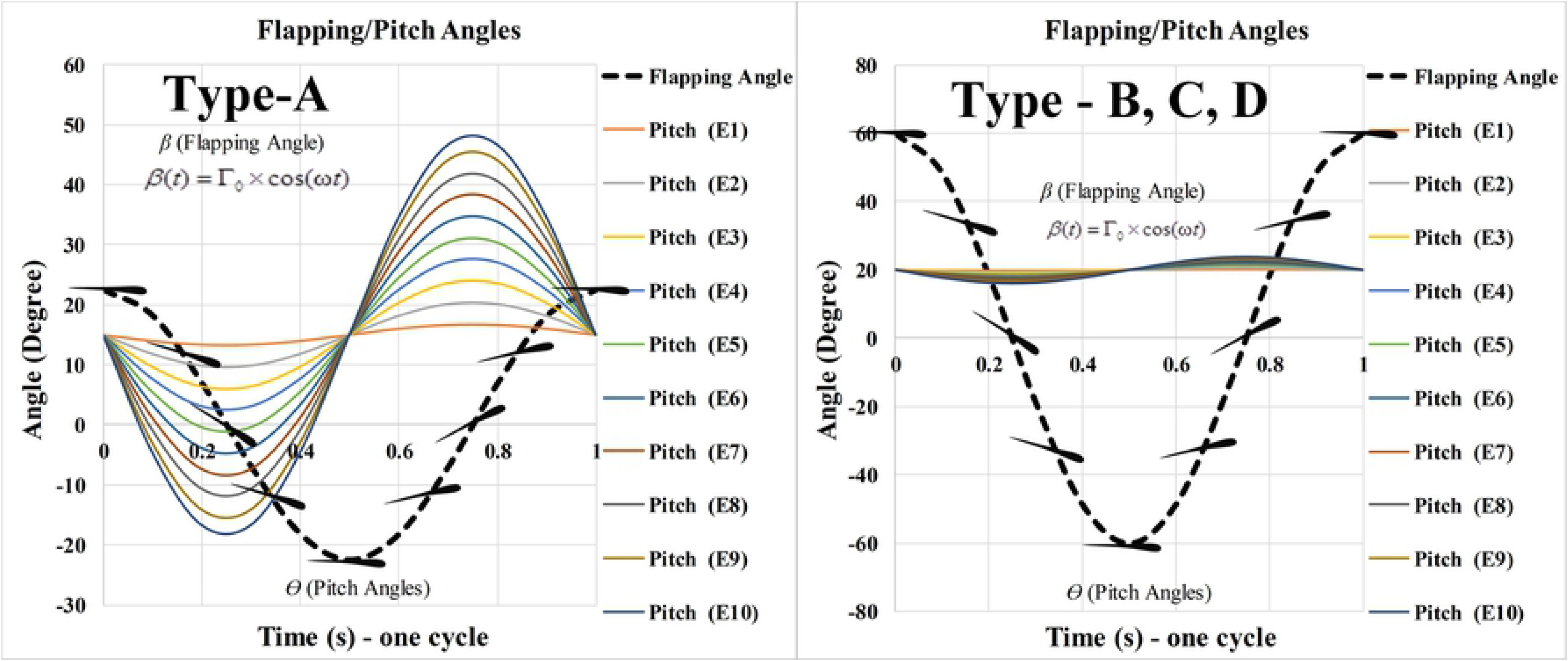
Variations of flapping and pitch angle in a cycle.

**S3 Fig.**
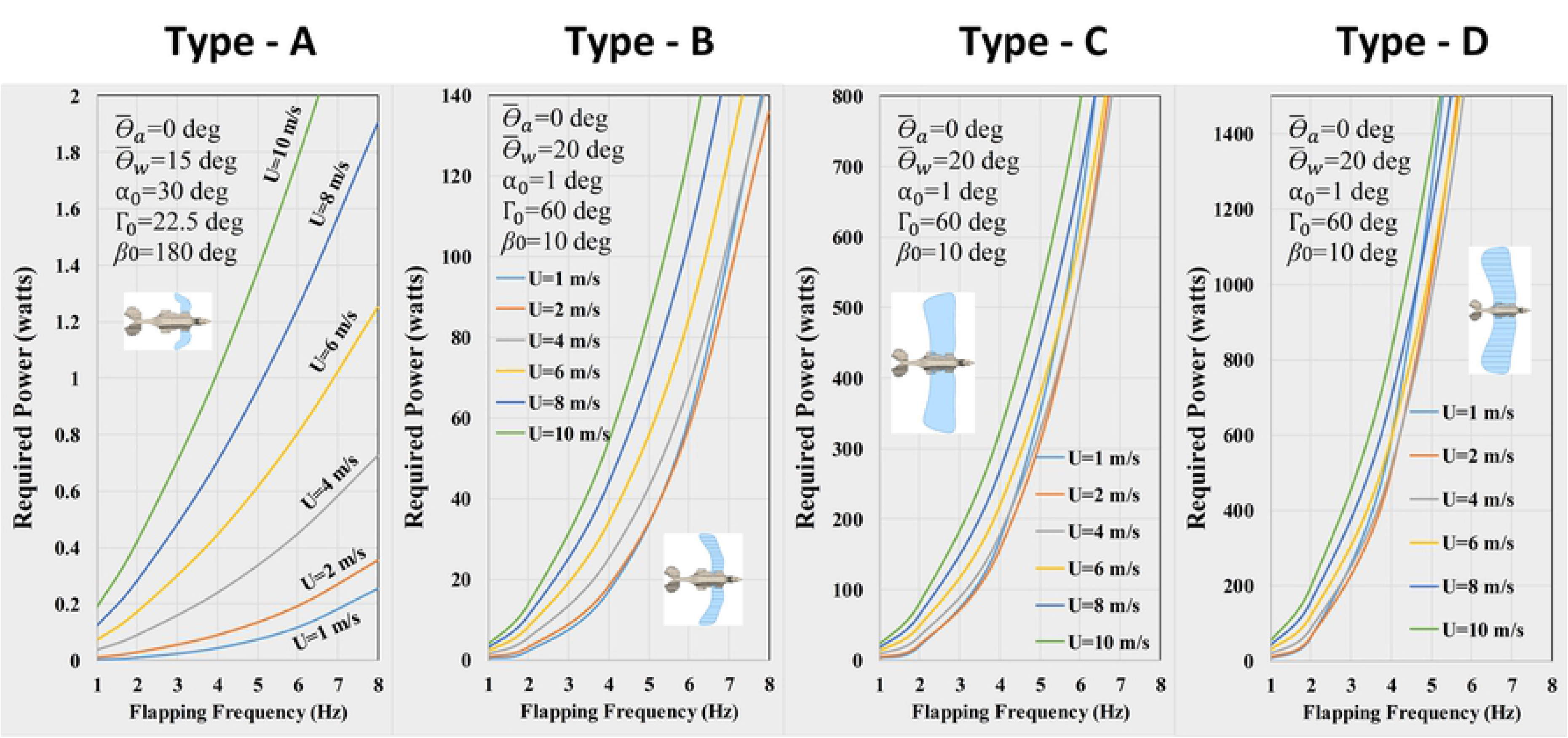
Variation in required metabolic power with flapping frequency at any velocit *Caudipteryx*.

**S4 Fig.**
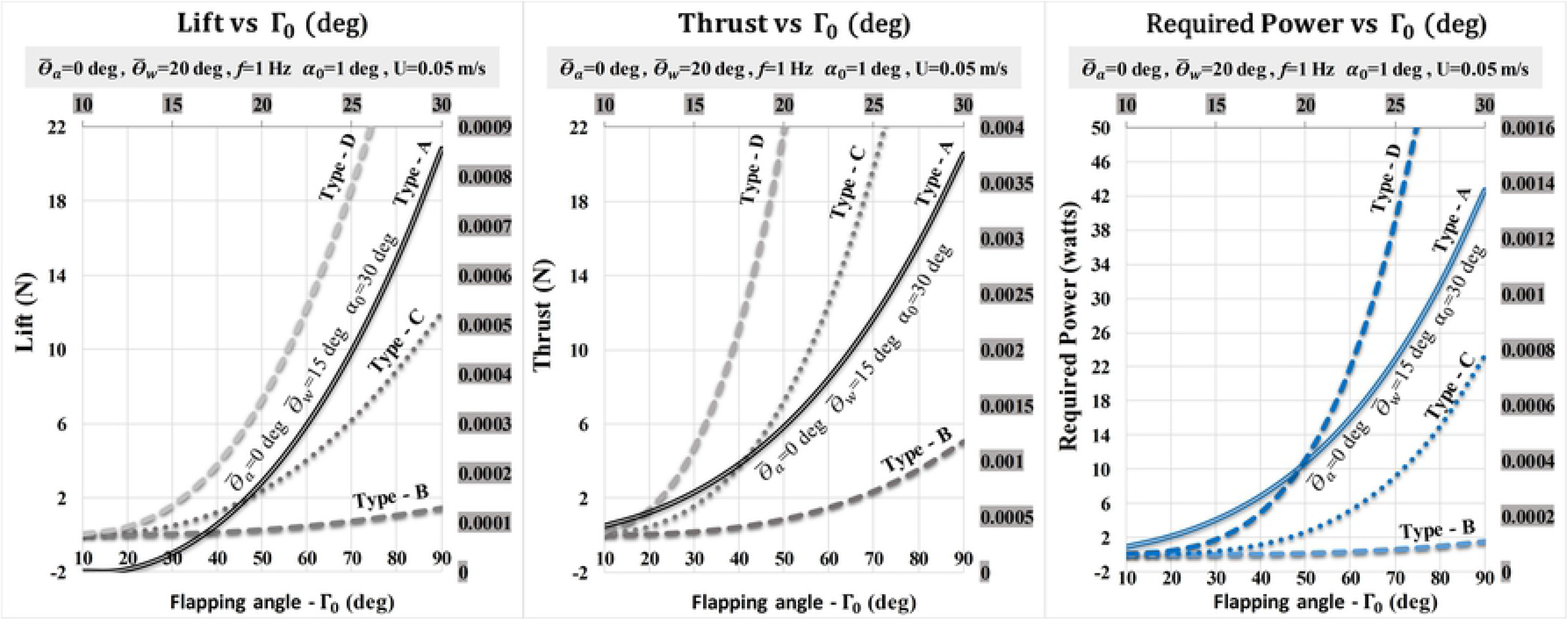
Variation in lift, thrust and power with flapping angle (Γ_0_) in a cycle when *U* is 0.05 m/s.

**S5 Fig.**
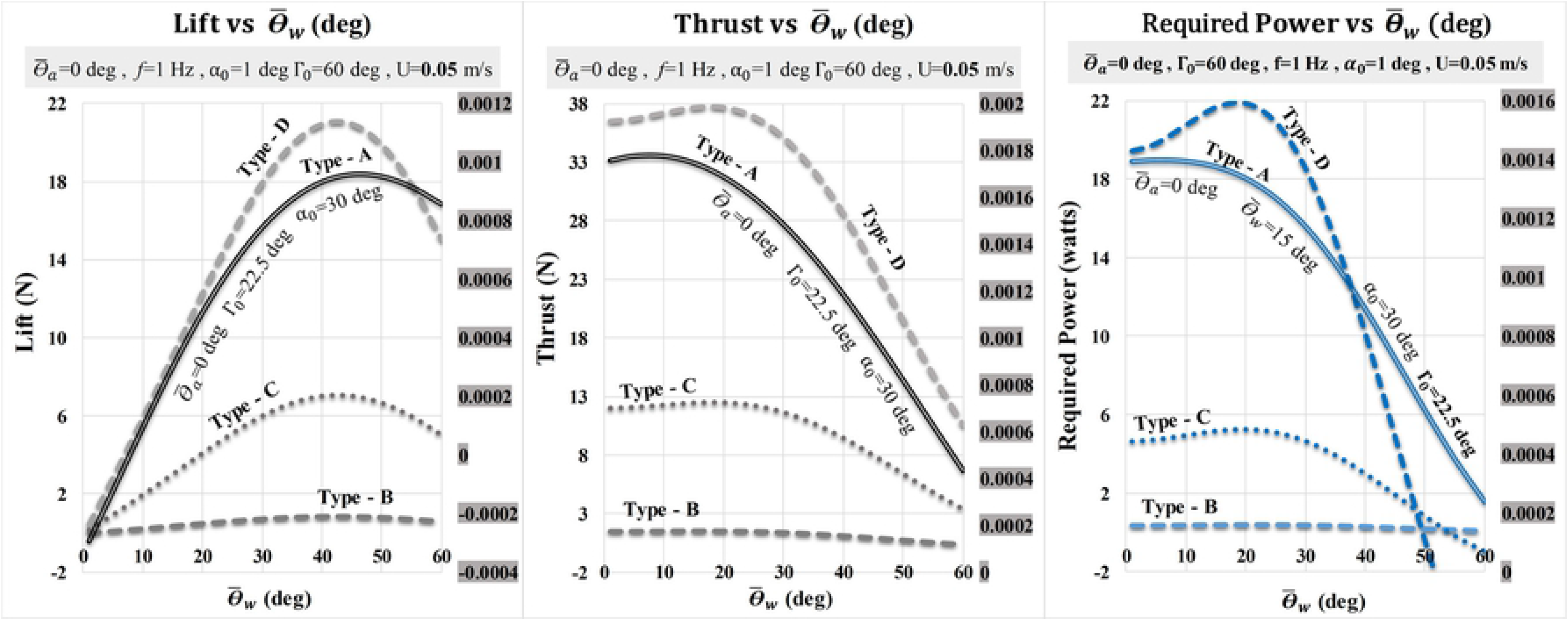
Variation in lift, thrust and power with 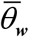 in a cycle when *U* is 0.05 m/s.

**S6 Fig.**
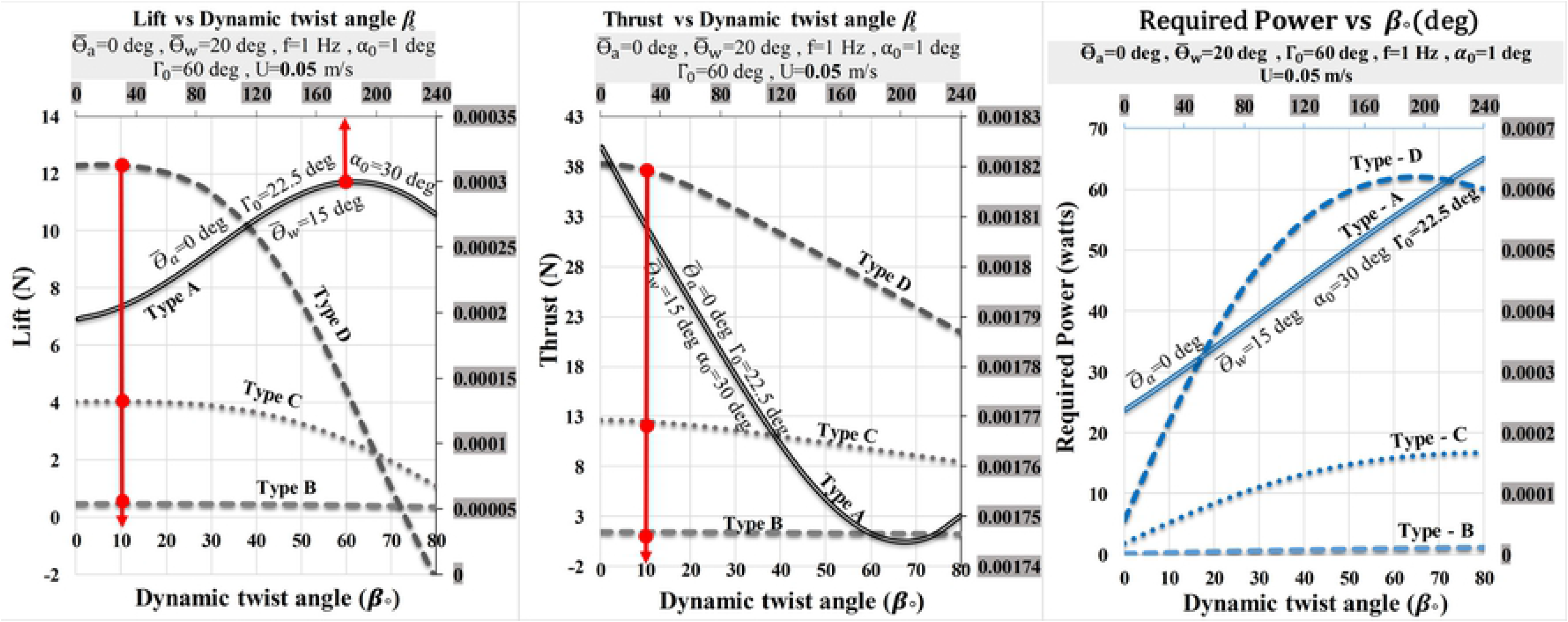
Variation in lift, thrust and power with dynamic twist angle (β_0_) in a cycle and when the velocity is almost zero.

**S7 Fig.**
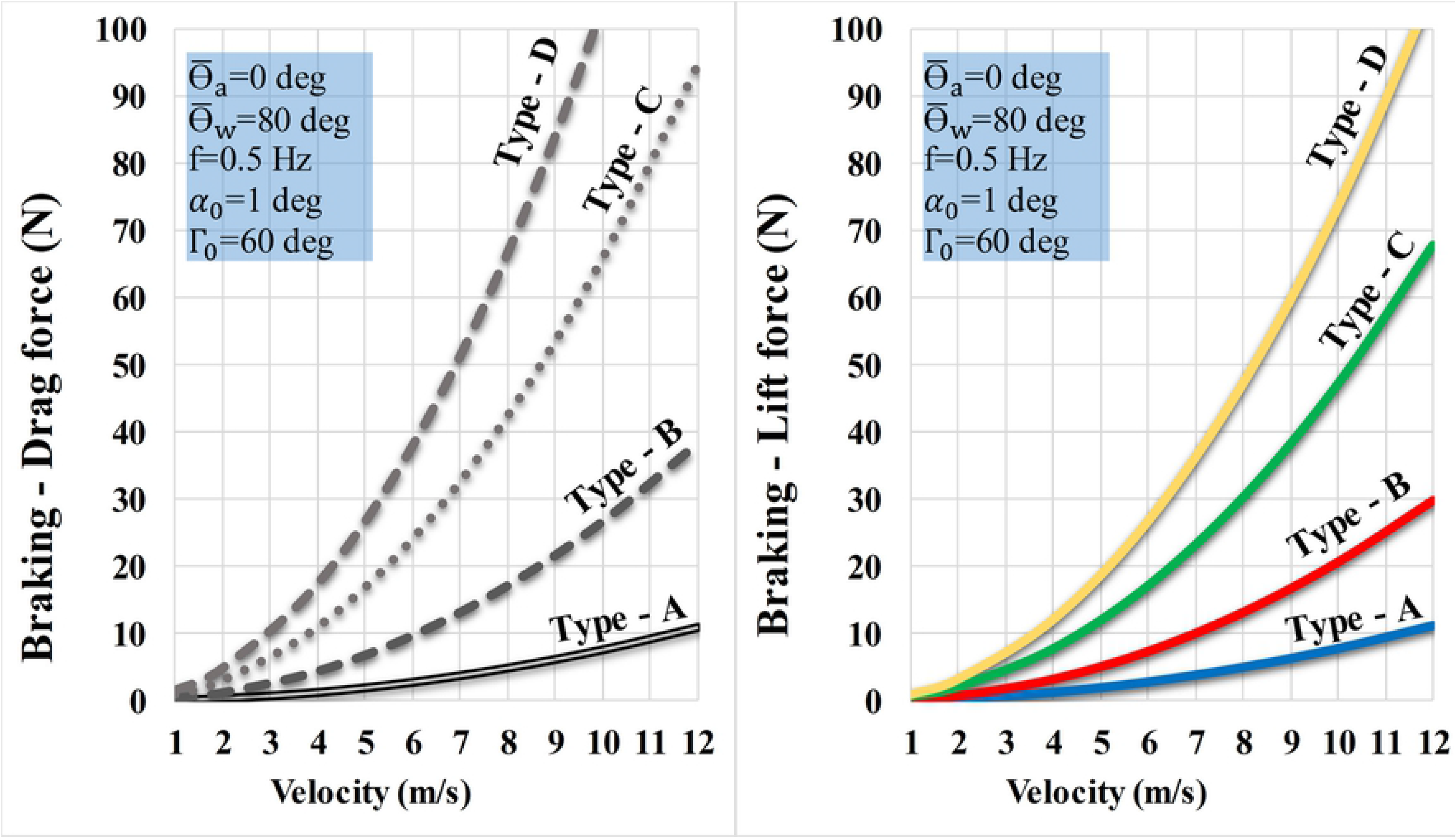
Variation in lift and drag with velocity in half of a cycle (downstroke) and high value of 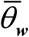.

**S8 Fig.**
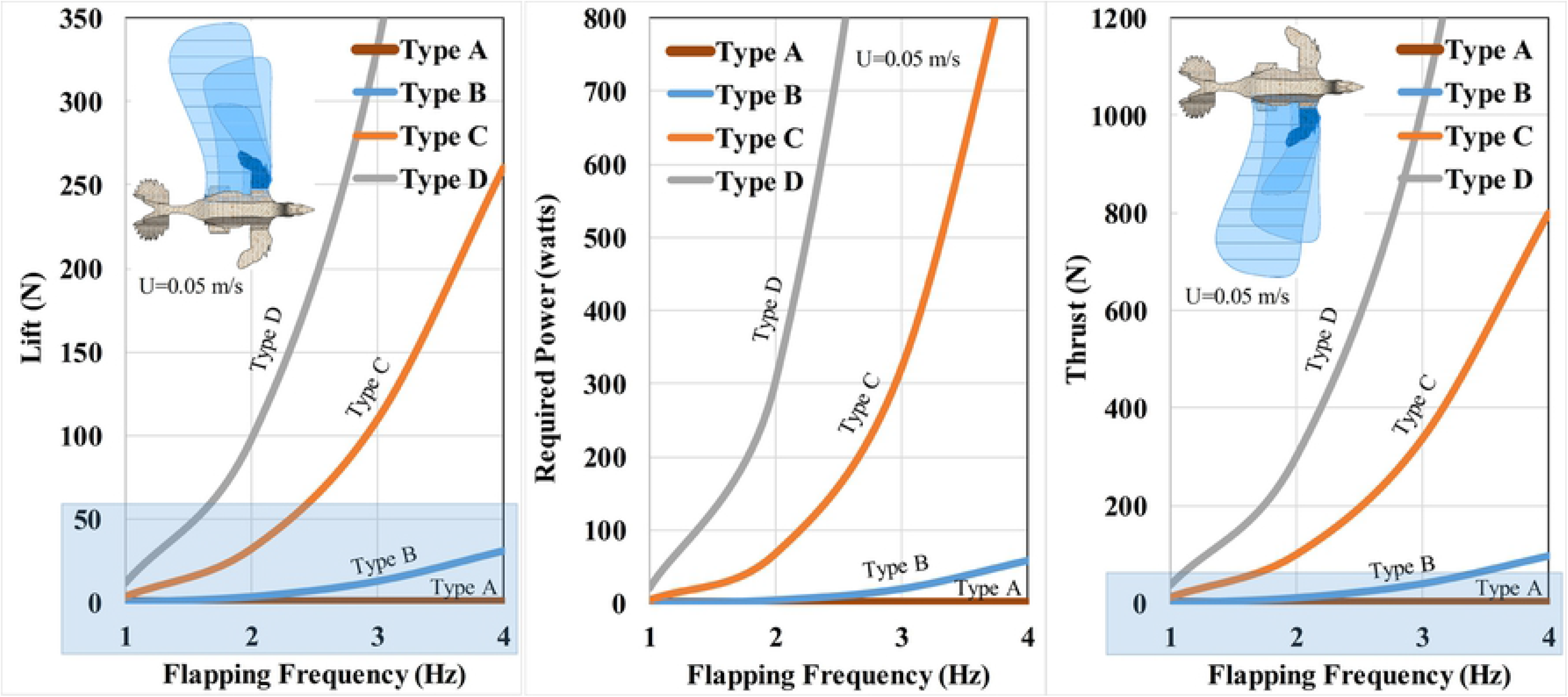
Variation in lift, thrust and required metabolic power with flapping frequency when the velocity is almost zero (*U*=0.05 m/s).

**S9 Fig.**
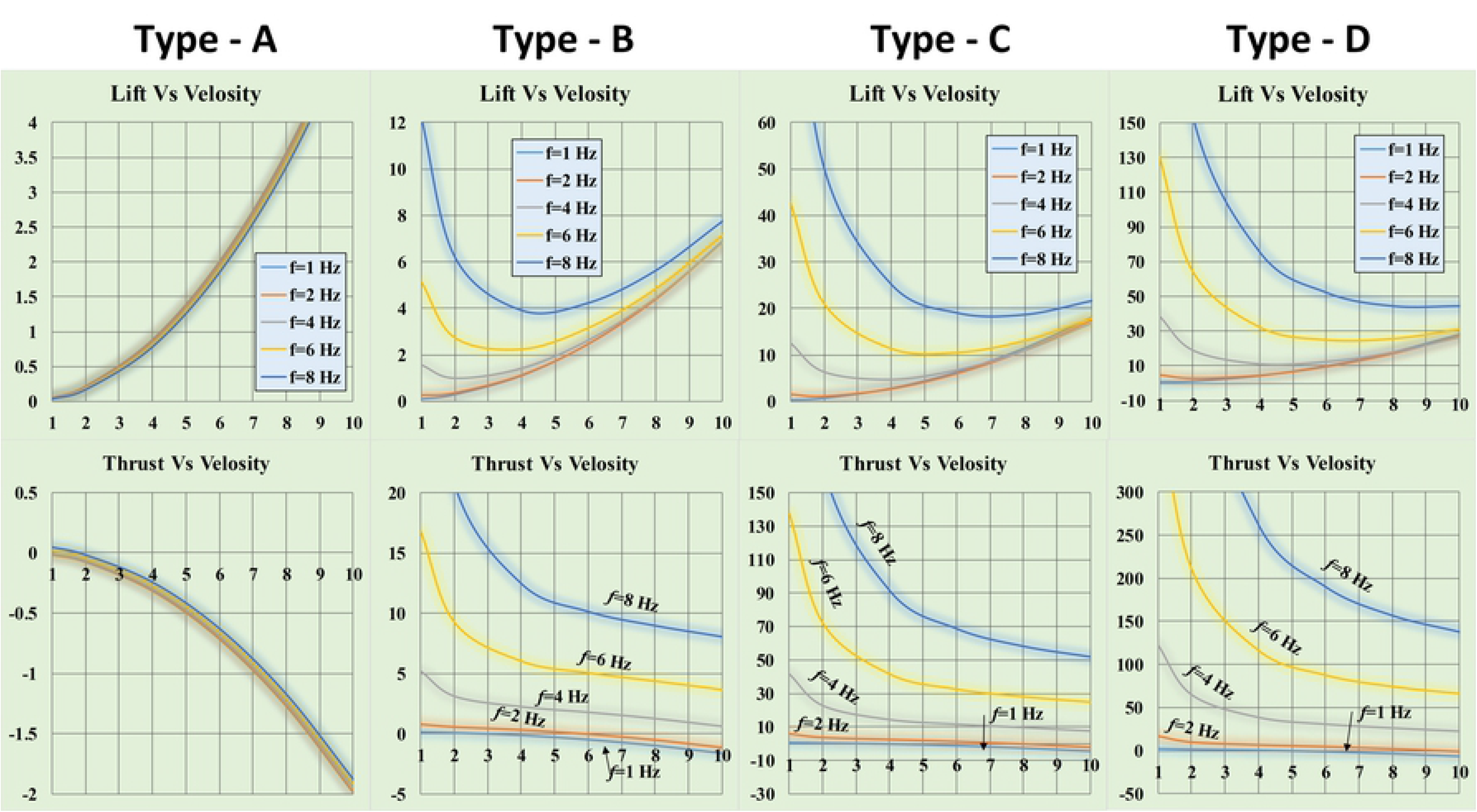
Variation in lift and thrust with velocity at any flapping frequency of *Caudipteryx*.

**S10 Fig.**
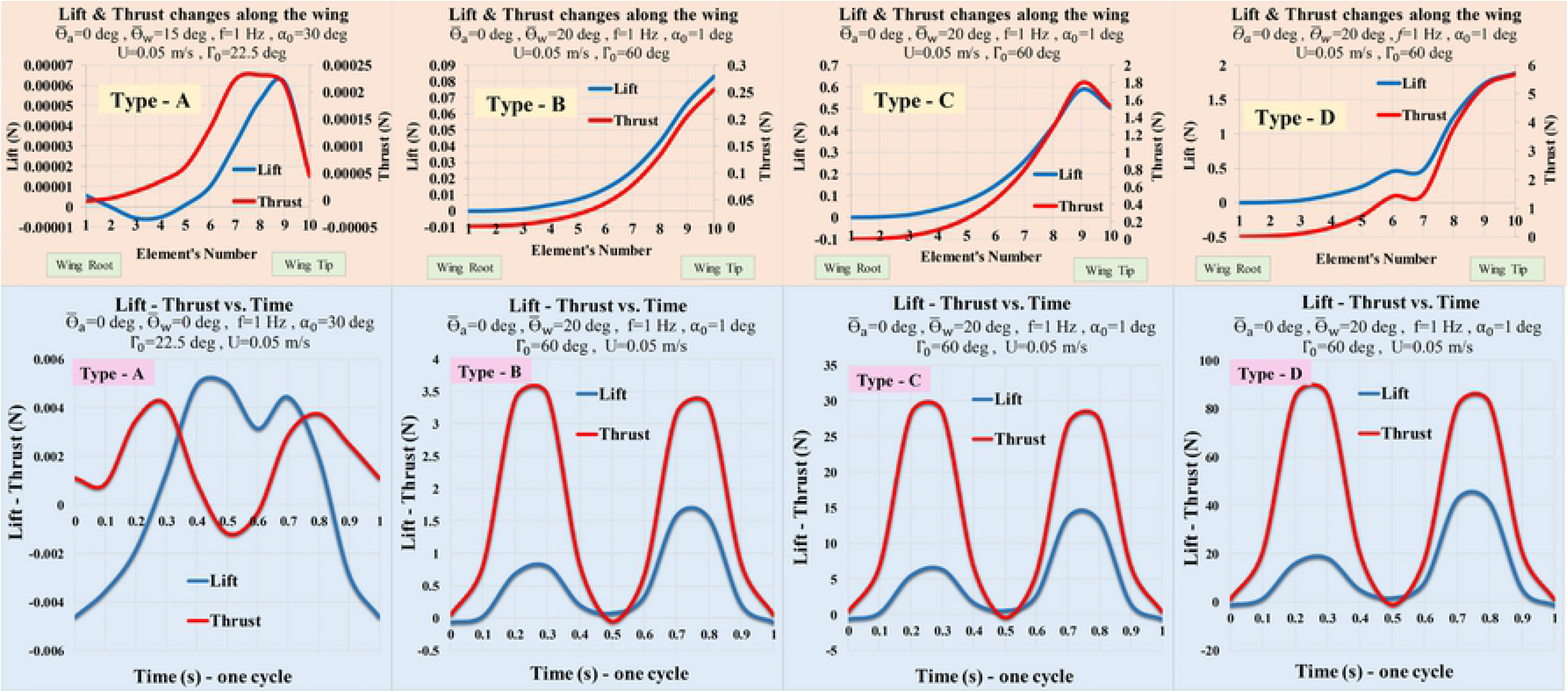
Lift and thrust changes along the wingspan. When the velocity is almost zero and in one cycle with the parameters shown in the Figure for types **A, B, C** and **D**, variations of lift and thrust forces in any element of wing from base to the tip are represented in top segment. Also the changes of lift and thrust in one cycle are illustrated in bottom segment of the Figure.

**S11 Fig.**
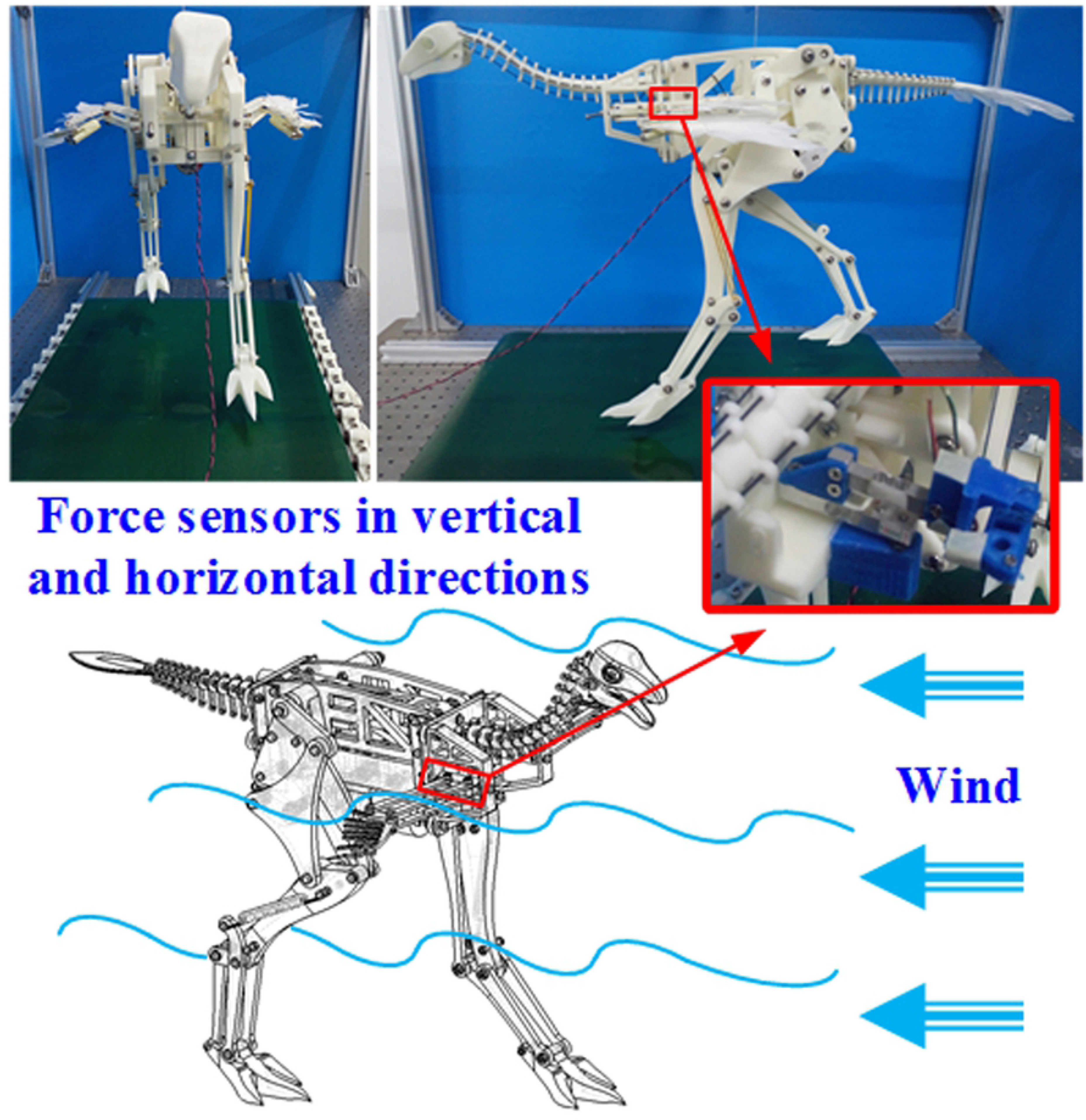
Reconstructed model (type A) of *Caudipteryx* on the test rig.

**S12 Fig.**
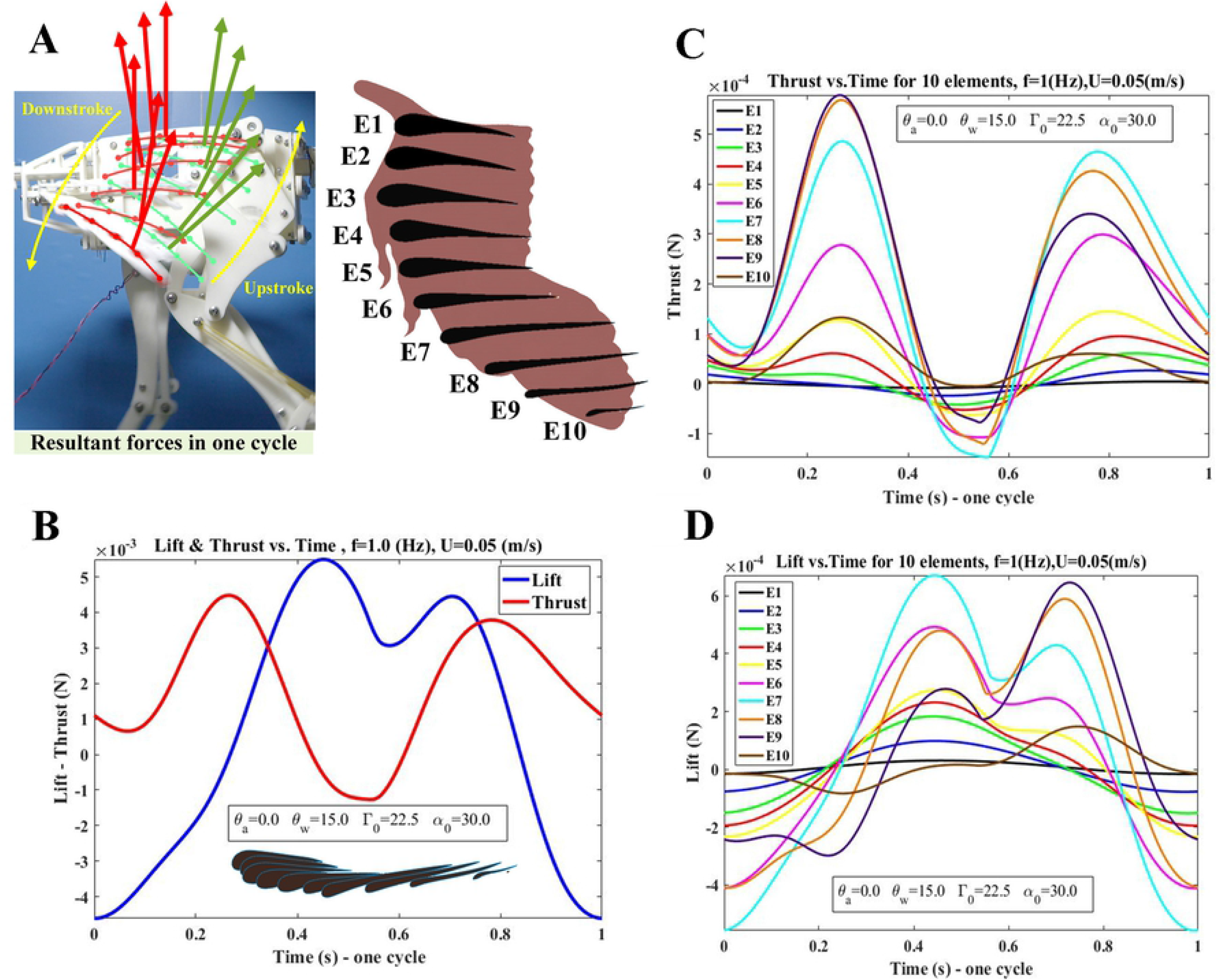
Lift and thrust forces of each element of wing of *Caudipteryx* (type **A**) (**A**), Resultant forces in one cycle. The wing of *Caudipteryx* is divided into ten elements along the wing span to better quantify the flight loads of each segment. (**B**), Variations of insignificant values of lift and thrust of whole wing in a cycle when the considered frequency is one. (**C**) and (**D**), Variations of insignificant values of thrust and lift of each segment of the wing to compare with each other’s. These values are computed for given parameters when the velocity of airflow is 0.05m/s and wing beat is equal to one in a cycle. It is obvious that by increasing flapping frequency the value of load increases.

**S13 Fig.**
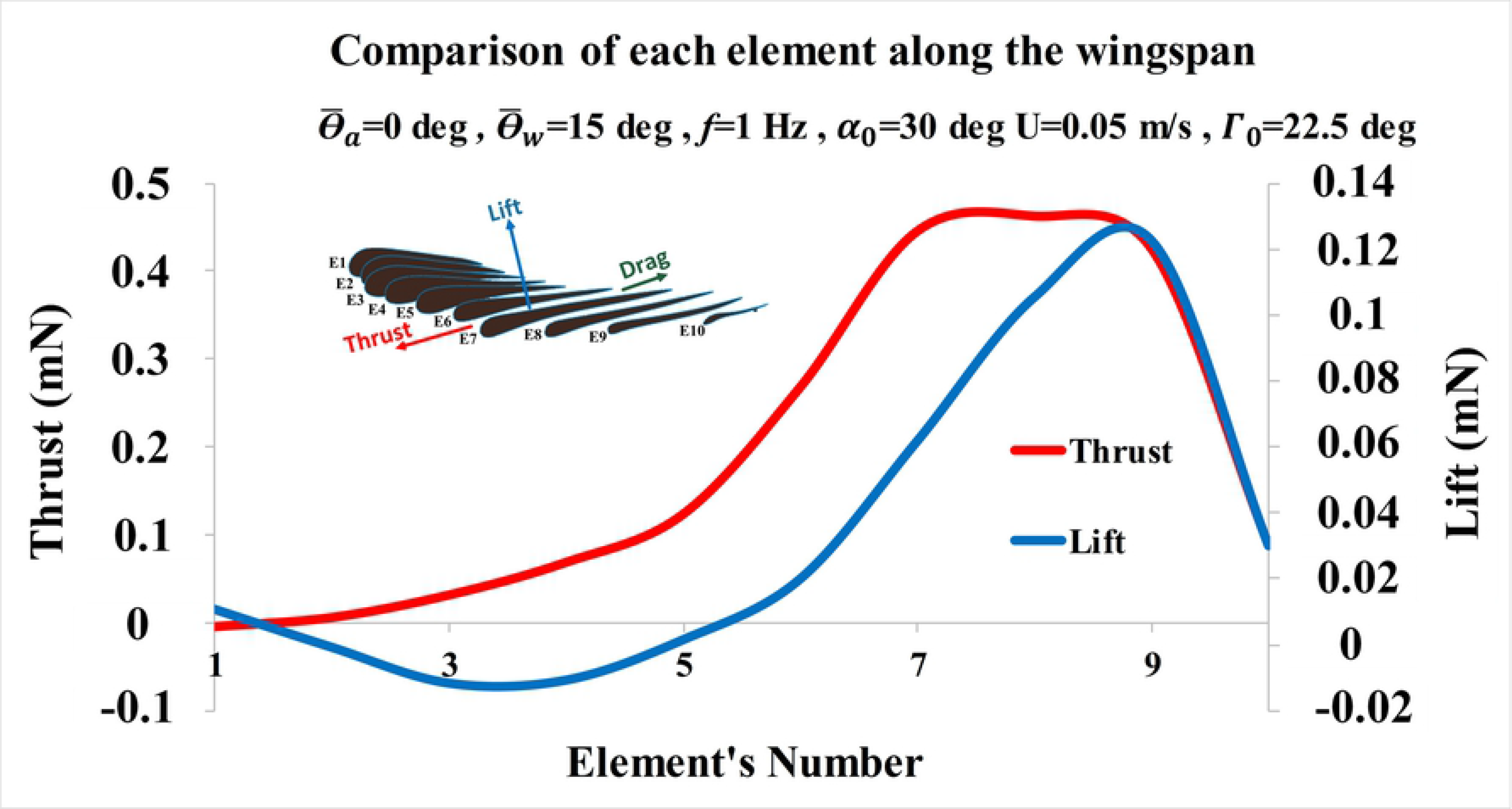
Comparisonof each element along the wingspan by measuring lift and thrust (type A) To compare any element along the wingspan of *Caudipteryx* and capture the properties of each segment, the insignificant values of lift and thrust of ten elements for the given parameters were deduced supposing the wing beat was one in a cycle. Lift from elements 4 to 9 and thrust from elements 2 to 9 considering distance to the wing root are meaningful but the wing tip (element 10) has insignificant value.

**S14 Fig.**
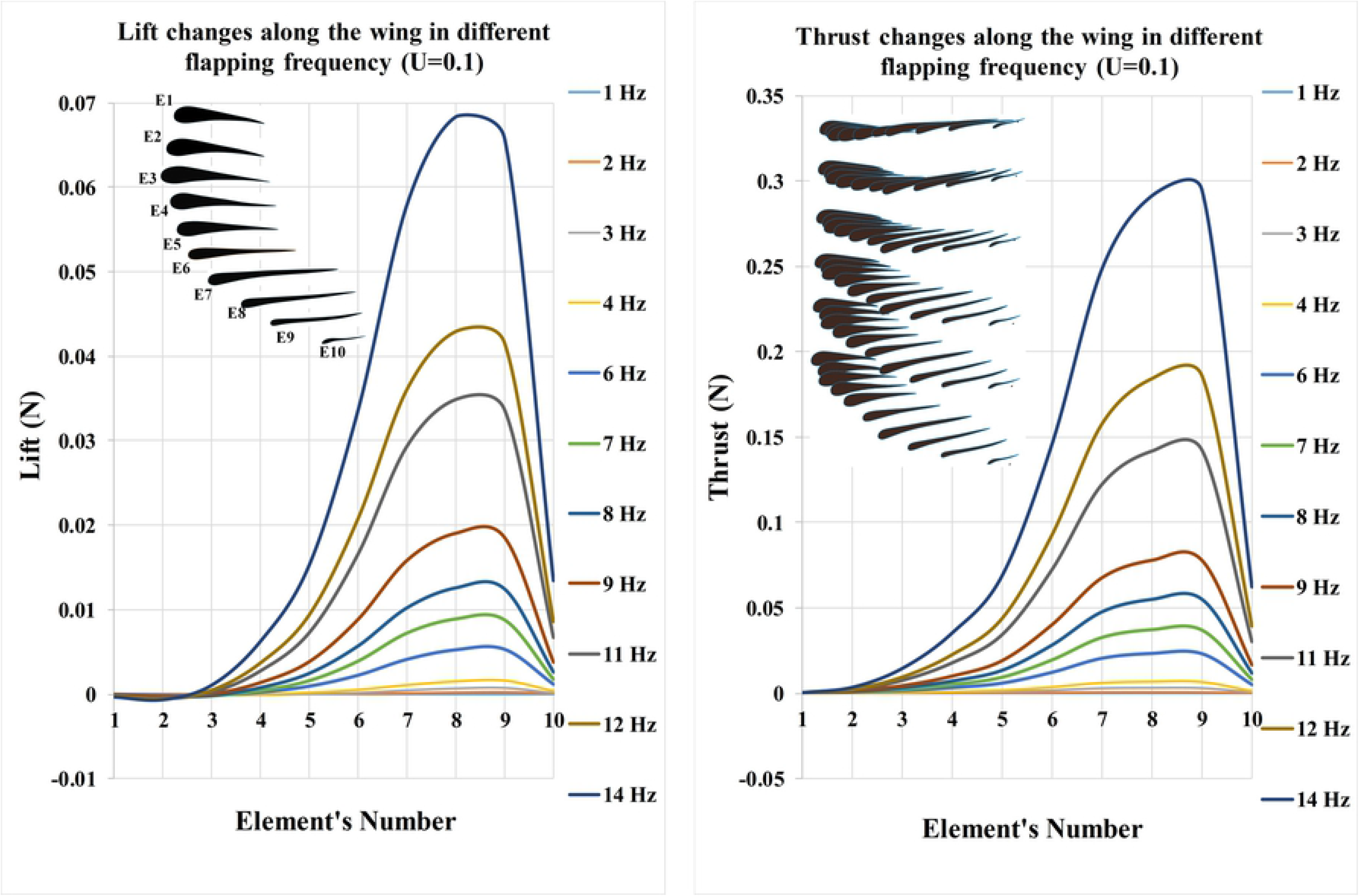
Comparisonof all elements along the wingspan of *Caudipteryx* at any wing beat by measuring lift and thrust (type A) Lift and thrust change from the base to the tip element by element at different flapping frequencies when the given parameters are 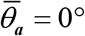, 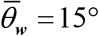, *α*_0_ = 30°, *β*_0_ = 180° and Γ_0_ = 22.5°. The significant values of lift and thrust began from somewhere ahead of the wing base between elements 3 and 4 to the wing tip (element 9) at any flapping frequency.

**S15 Fig.**
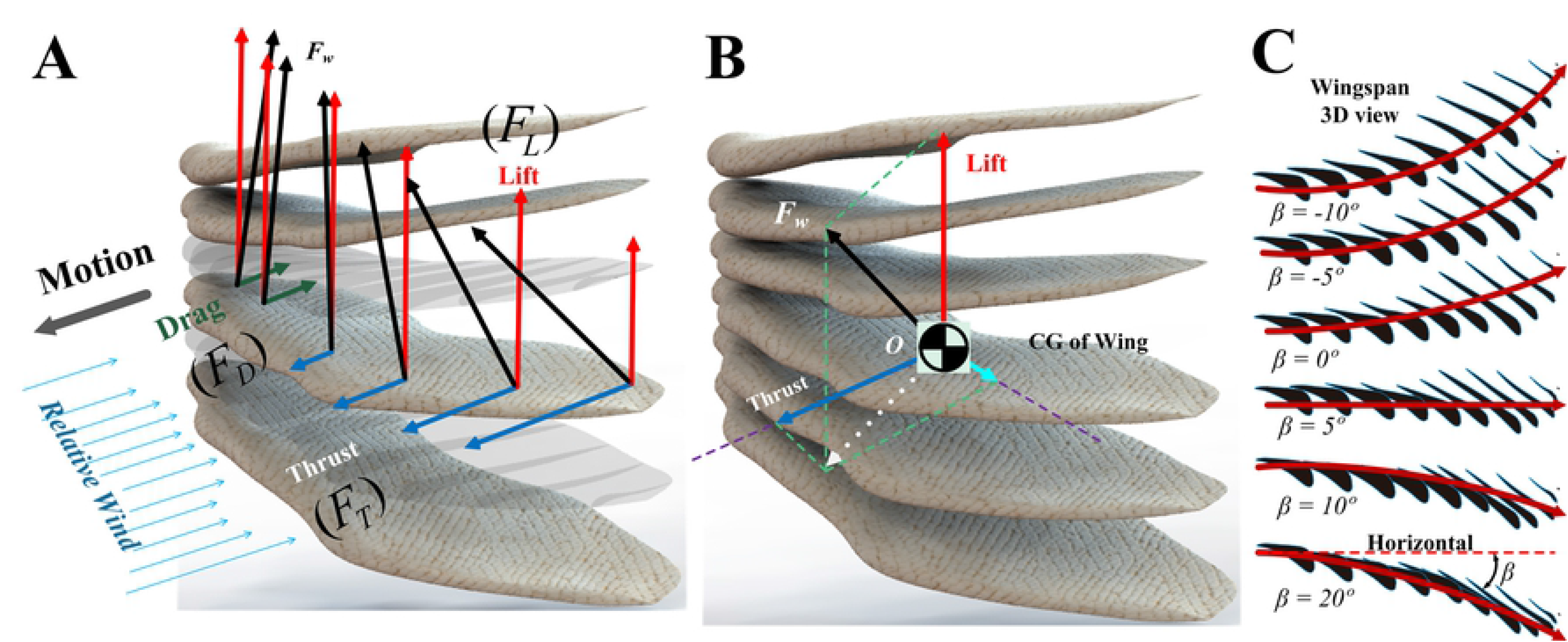
Influence of aerodynamic forces on unfolded wing of *Caudipteryx* at some fixed flapping angles during running (this process can happen in the downstroke) (**A**) and (**B**), ***F**_w_* is the transverse force composed of three components of thrust/drag in motion, lift in vertical and a force along the wingspan direction. (**C**), Wing of *Caudipteryx* is supposed to be fixed in six flapping angles (***β*** is equal to -10, -5, 0, 5, 10, 20 degrees) and twisted along the wing span similar to modern birds.

**S16 Fig.**
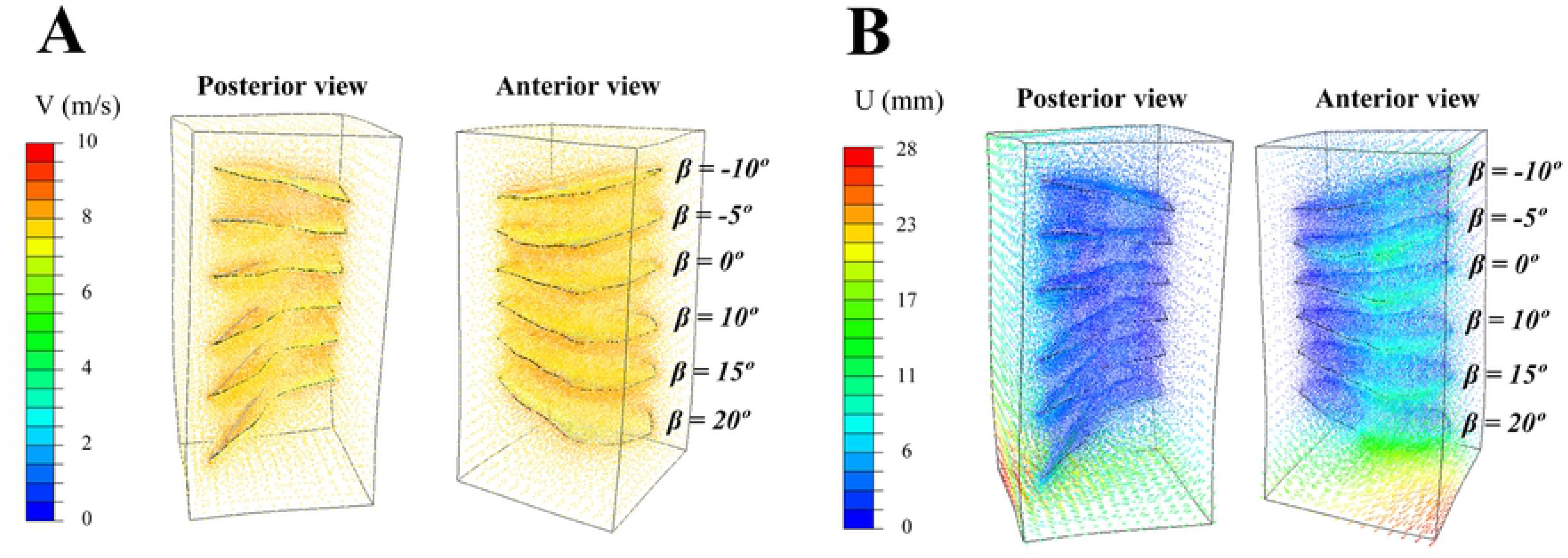
Airflow around the wing. (**A**), in order to simulate all states in the same condition, all cases are compared and arranged together. Hence, airflow goes through all states of the wing of *Caudipteryx* at the velocity of 8 m/s, it is shown around the wing. (**B**), Displacement (in meters) of the nodes of airflow while passing the wing.

**S17 Fig.**
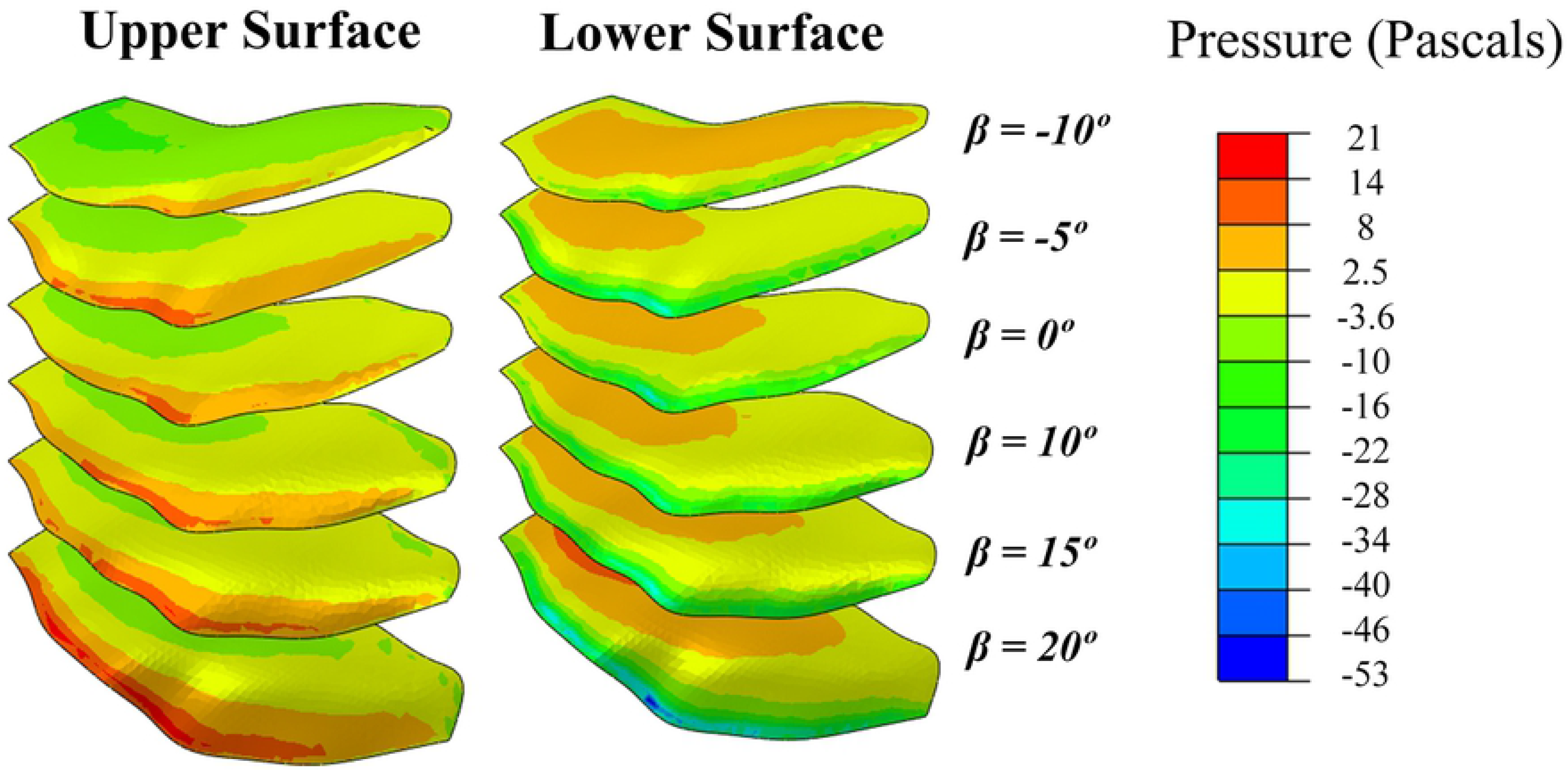
Airflow produces pressure (in Pascals) on the top and bottom surfaces of the wing. In bottom surface, the value of pressure is higher than that in the top surface. This creates lift force. Therefore, for different flapping angle, the gradient of pressure must be positive to generate lift.

**S18 Fig.**
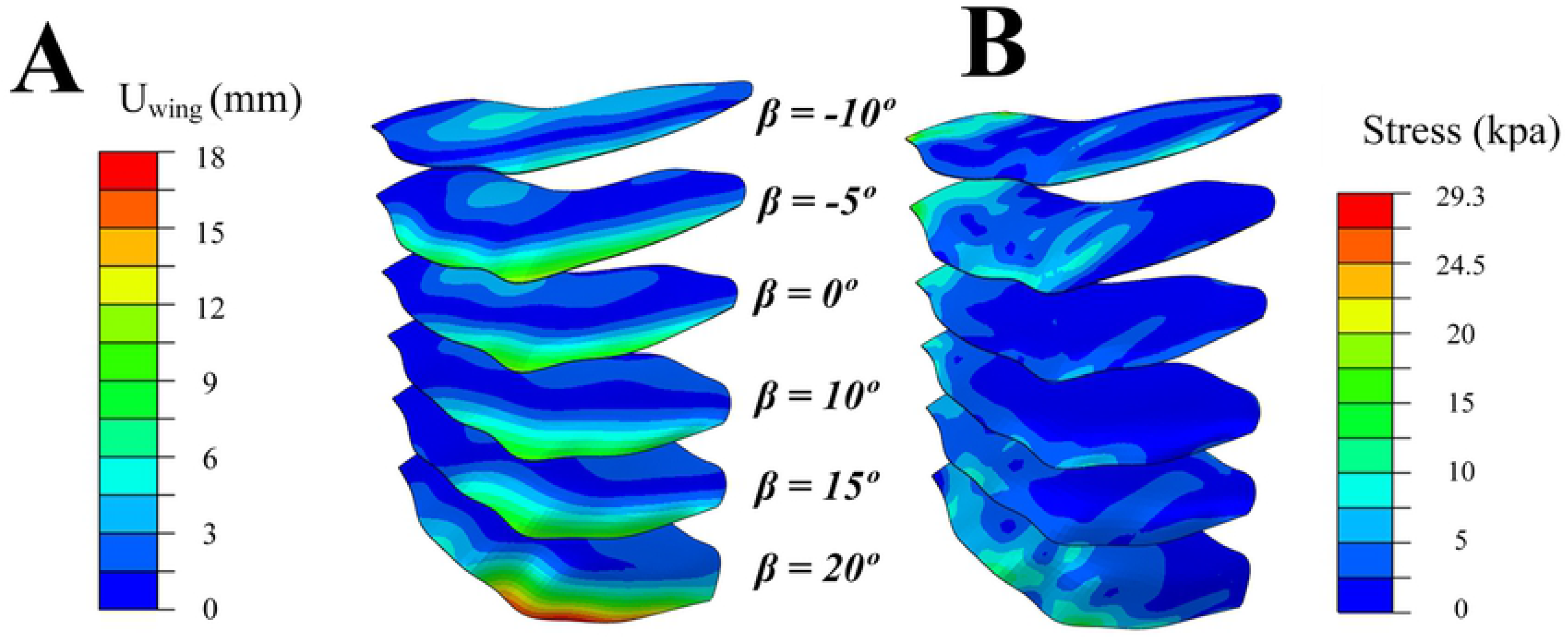
Displacement and stress of a wing. (**A**), Displacement of the wing of *Caudipteryx* in meters. The wing lead has most deflection for any flapping angle. (**B**), Stress on the wing of *Caudipteryx* under the effect of airflow in Pascals (the speed of airflow is 8 m/s). The wing base (shoulder joint) and forelimb skeleton bear most bending and torsional stresses.

**S19 Fig.**
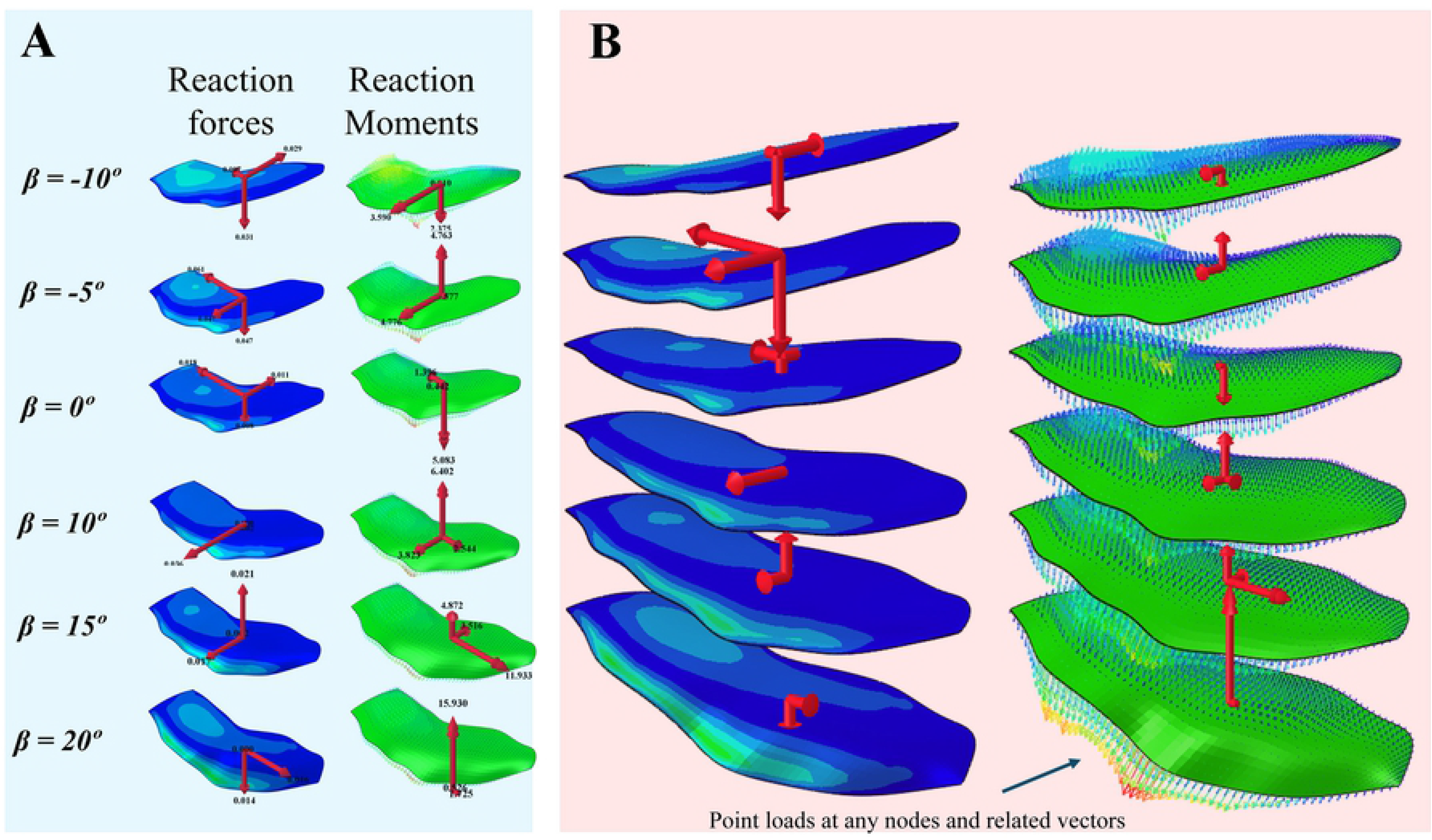
Reaction forces and moments of the wing of *Caudipteryx*. (**A**), the reaction forces and moments are illustrated (the *Caudipteryx*’s velocity is 8 m/s in each state). (**B**), Point loads (N) at any nodes and related vectors. The average of transverse force in all cases are composed of three components, thrust/drag force in motion, lift in vertical and a force along the wingspan direction. It illustrates that thrust can appear in some flapping angles and the aerodynamic force in wingspan direction is useful to expand the wing.

**S20 Fig.**
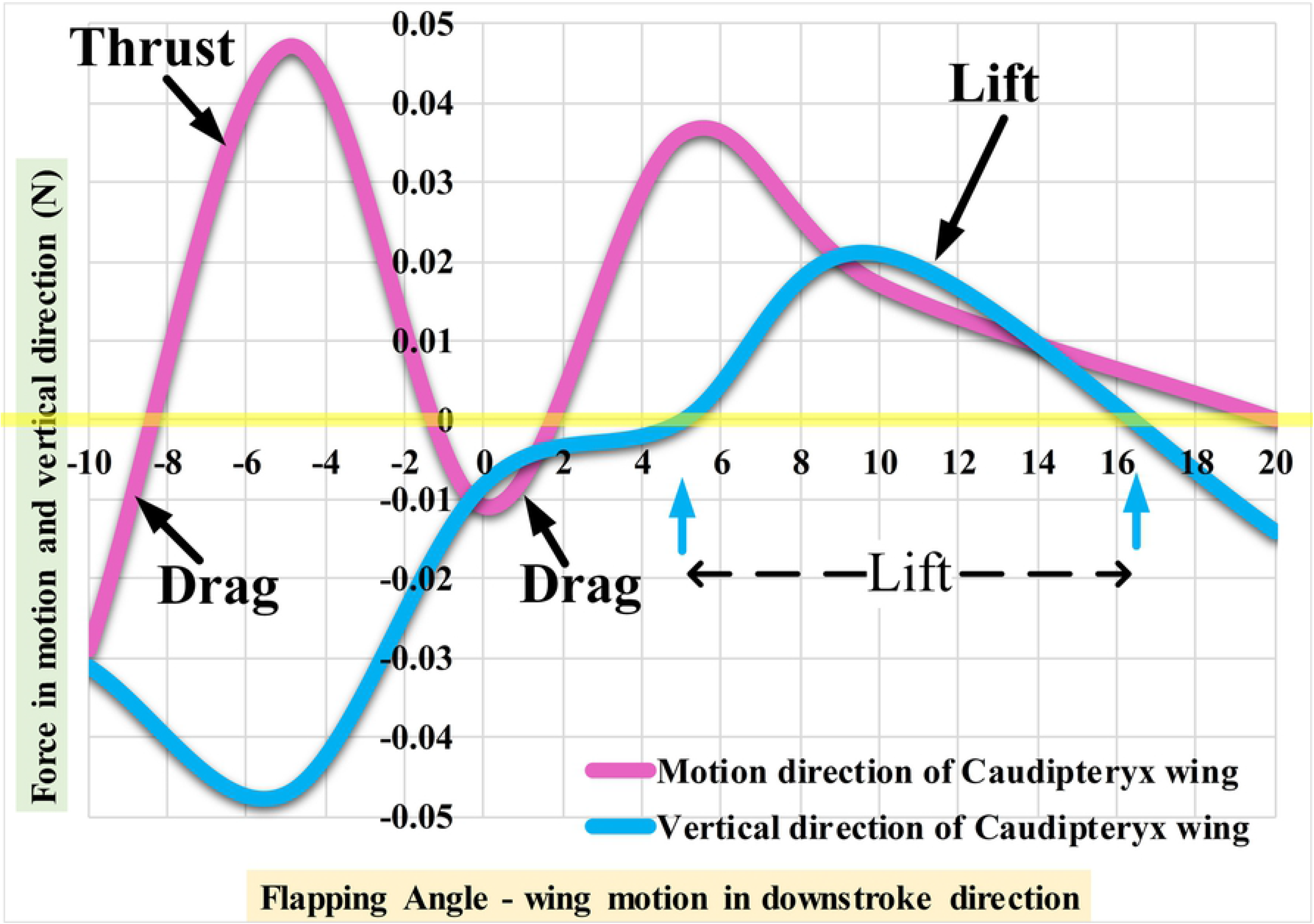
The force in horizontal and vertical directions versus the flapping angle from -10 to 20 deg. The average lift and thrust/drag forces generated by simulated wing of *Caudipteryx* under the effect of airflow are different in various cases. It depends on the wingspan, flapping and twist angles of the wing and the general thickness of the wing along the span (each section/airfoil has its own properties such as angle of attack along the wing). For the sake of similarity to downstroke process, angles of attack along the wingspan are assigned in such a way that the possibility of thrust force becomes more than that of drag. Hence, from the flapping angle of -10 to 20 degrees, the possibility of having thrust force is more than that of drag. In addition, in this case, the possible lift force occurs from 5 degrees to 16 degrees for the flapping angles. Namely, only in these angles, the wing has lift and thrust at the same time. Therefore, *Caudipteryx* could obtain lift and thrust if it adjusted its wings in a proper flapping and pitching angles by using its unfolded wings during running fast. It is obvious that *Caudipteryx* could adjust its wings to get lift and drag at the same time.

**S21 Fig.**
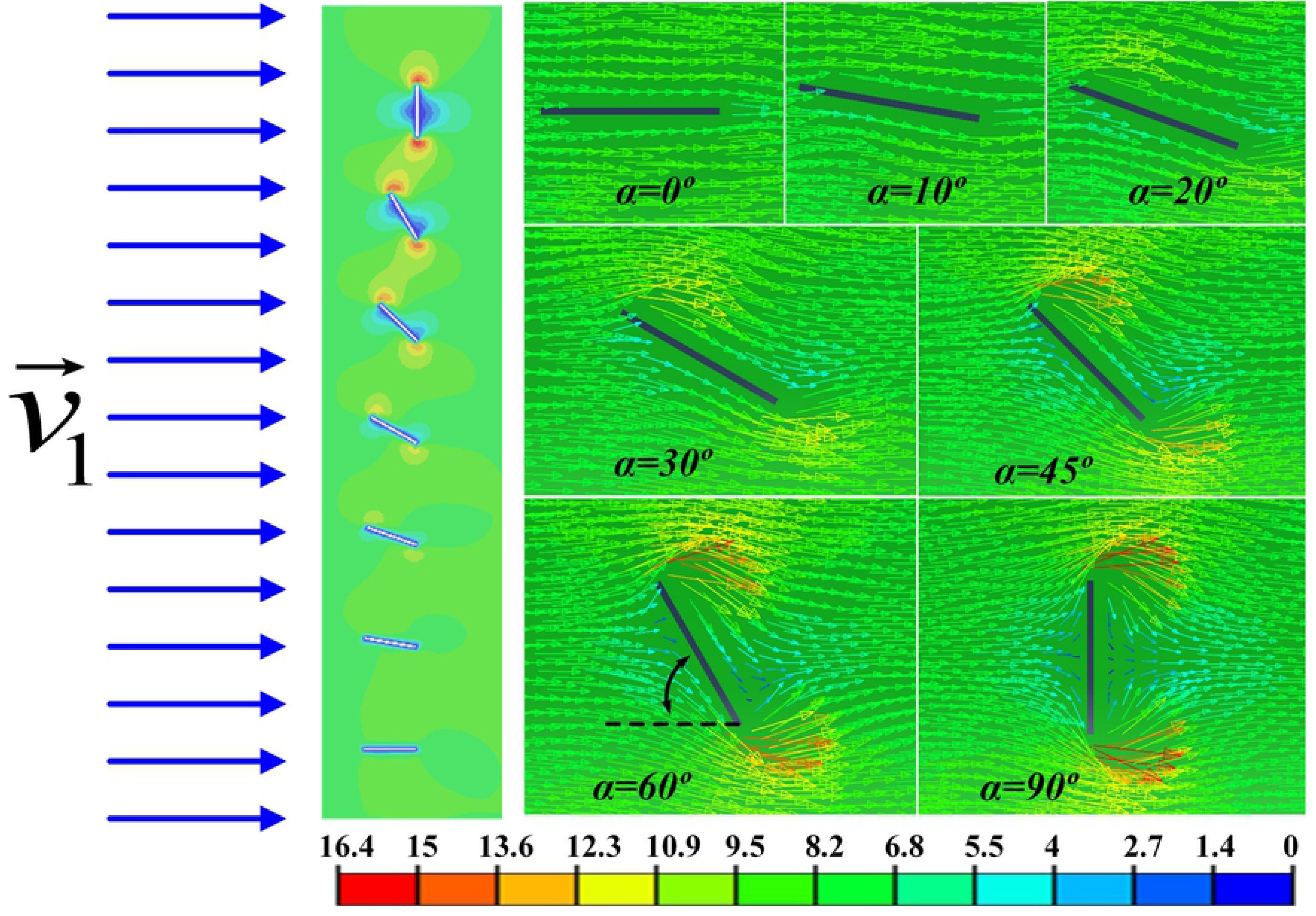
Simulation of simplified model of rectangular wing. Velocity vectors of airflow around the rectangular plate (both bottom surface and upper surface of the rectangular wing) in various angles of attack (0° to 90°) under the effect of initial velocity of ***v***_1_ = 8 m/s. The stalling occurs after 45 degrees, hence, drag force increases. The maximum speed of the airflow around the plate is 16 m/s (twice of inlet speed) near the edges.

**S1 Table.** Comparison between different modern birds in self-weight, wing area, wing loading *(W* / *S*), wing span and velocity [4]. The values of velocity are calculated from equation (2). In general, larger birds have to fly faster.

**S2 Table.** Lift and thrust forces of reconstructed robot of *Caudipteryx* on the test rig (Type A) at different velocities.

**S3 Table.** Comparison between modern birds in body mas, wing area, flap angle, wing beat and velocity.

**S4 Table.** Comparison between flightless dinosaurs in body mass, wing area, flap angle, wing span and wing loading.

## Data Availability Statement

All relevant data are within the paper and its Supporting Information files.

## References

1. Ostrom, J.H. Bird flight: how did it begin? Am. Sci. 67, 46–56 (1979).

2. Forster, C.A., Sampson, S.D., Chiappe, L.M. & Krause, D.W. The theropod ancestry of birds: new evidence from the late cretaceous of madagascar. Science 279, 1915–1919 (1998).

3. Garner, J.P., Taylor, G.K. & Thomas, A.L.R. On the origins of birds: the sequence of character acquisition in the evolution of avian flight. Proc. R. Soc. Lond. B. 26, 1259–1266 (1999).

4. Sereno, P.C. The evolution of dinosaurs. Science 284, 2137–2147 (1999).

5. Lewin, R. How did vertebrates take to the air? Science New Series 221, 38–39 (1983).

6. Gibbons, A. New feathered fossil brings dinosaurs and birds closer. Science New Series 274, 720–721 (1996).

7. Prum, R.O. Dinosaurs take to the air. Nature 421, 323–324 (2003).

8. Henderson, D.M. Estimating the masses and centers of mass of extinct animals by 3-d mathematical slicing. Paleobiology 25, 88–106 (1999).

9. Ji, S.A., Ji, Q. & Padian, K. Biostratigraphy of new pterosaurs from china. Nature 398, 573–574 (1999).

10. Dyke, G.J. & Norell, M.A. Caudipteryx as a non avialan theropod rather than a flightless bird. Acta Palaeontol Pol. 50(1), 101–116 (2005).

11. Padian, K. When is a bird not a bird? Nature 393, 729–730 (1998).

12. Qiang, J., Currie, P.J., Norell, M.A. & Shuan, J. Two feathered dinosaurs from northeastern china. Nature 393, 753–761 (1998).

13. Swisher, C.C., Wang, Y.Q., Wang, X.L. & Xu, X. Cretaceous age for the feathered dinosaurs of liaoning, china. Nature 400, 58–61 (1999).

14. Xu, X., Wang, X.L. & Wu, X.C. A dromaeosaurid dinosaur with a filamentous integument from the yixian formation of china. Nature 401, 262–266 (1999).

15. Zhou, Z. The origin and early evolution of birds: discoveries, disputes, and perspectives from fossil evidence. Naturwissenschaften 91, 455–471 (2004).

16. Xu, X. et al. Four-winged dinosaurs from china. Nature 421, 335–340 (2003).

17. Zhou, Z. & Zhang, F.C. Origin and early evolution of feathers: evidence from the early cretaceous of china. Acta Zoologica Sinica 125–128 (2006).

18. Zhou, Z. & Wang, X. A new species of Caudipteryx from the yixian formation of liaoning, northeast china. Vertebr. PalAsiat 38, 104–122 (2000).

19. Zhou, Z., Barrett, P.M. & Hilton, J. An exceptionally preserved lower cretaceous ecosystem. Nature 421, 807–814 (2003).

20. Jones, T.D., Farlow, J.O., Ruben, J.A, Henderson, D.M. & Hillenius, W.J. Cursoriality in bipedal archosaurs. Nature 406, 716–718 (2000).

21. Norell, M., Ji, Q., Gao, K., Yuan, C., Zhao, Y. & Wang, L. Modern feathers on a non-volant dinosaur. Nature 416, 36–37 (2002).

22. Lee, M.S.Y., Cau, A., Naish, D. & Dyke, G.J. Sustained miniaturization and anatomical innovation in the dinosaurian ancestors of birds. Science 345, 562–566 (2014).

23. Zhou, Z. Evolutionary radiation of the jehol biota: chronological and ecological perspectives. Geol. J. 41, 377–393 (2006).

24. Zhou, Z., Wang, X.L., Zhang, F.C. & Xu, X. Important features of Caudipteryx evidence from two nearly complete new specimens. Vertebr PalAsiat 38, 241–254 (2000).

25. Norberg, U.M. VERTEBRATE FLIGHT IN ZOOPHYSIOLOGY. p. ISBN: 9783642838507 (1989).

26. Delaurier, J.D. & Larijani, R.F. A Nonlinear Aeroelastic Model for the Study of Flapping Wing Flight. pp. Chapter 18, Published by the American Institute of Aeronautics and Astronautics (2001).

27. Delaurier, J.D. An aerodynamic model for flapping wing flight. Aeronautical journal 97, 125–130 (1993).

28. Delaurier, J.D. The development of an efficient ornithopter wing. Aeronautical journal 97, 153–162 (1993).

29. Delaurier, J.D. An ornithopter wing design. Canadian Aeronautics and space Journal 40, 10–18 (1994).

30. Delaurier, J.D. Drag of wings with cambered airfoils and partial leading edge suction. Journal of Aircraft 20, 882–886 (1983).

31. Delaurier, J.D. The development and testing of a full-scale piloted ornithopter. Canadian Aeronautics and Space Journal 45, 72–82 (1999).

32. Delaurier, J.D. & Harris J.M. A study of mechanical flapping wing flight. Aeronautical Journal 97, 277–286 (1993).

33. Jones, R.T. The Unsteady Lift of a Wing of Finite Aspect Ratio.p. NACA Report 681 (1940).

34. Hoerner, S.F. Fluid dynamic drag. Brick Town, NJ 2, 1–16 (1965).

35. Garrick, I.E. Propulsion of a Flapping and Oscillating Aerofoil.p. NACA Report 567 (1936).

36. Hoerner, S.F. Pressure drag, fluid dynamic drag. Brick Town, NJ, 3–16 (1965).

37. Christiansen, P. & Farina, R.A. Mass prediction in theropod dinosaurs. Historical Biology. 16(2-4), 85–92 (2004).

38. Hutchinson, J.R. & Allen, V. The evolutionary continuum of limb function from early theropods to birds. Naturwissenschaften. DOI 10.1007/s00114-008-0488-3 (2008).

39. Robertson, A.M.B & Biewener, A.A. Muscle function during takeoff and landing flight in the pigeon (columba livia). The Journal of Experimental Biology 215, 4104–4114 (2012).

40. Benton, M.J. VERTEBRATE PALEONTOLOGY. Third edition 2005, Blackwell Science Ltd (2005).

41. Kovacs, C.E. & Meyers, R.A. Anatomy and histochemistry of flight muscles in a wing-propelled diving bird, the atlantic puffin, fratercula arctica. journal of morphology 244, 109–125 (2000).

